# NAC61 regulates late-and post-ripening associated processes in grapes involving a NAC60-dependent regulatory network

**DOI:** 10.1101/2023.05.17.541132

**Authors:** Chiara Foresti, Luis Orduña, José Tomás Matus, Elodie Vandelle, Davide Danzi, Oscar Bellon, Giovanni Battista Tornielli, Alessandra Amato, Sara Zenoni

## Abstract

During late-and post-ripening stages, grape berry undergoes profound biochemical and physiological changes whose molecular control is poorly understood. Here, we report the role of NAC61, a grapevine NAC transcription factor, in regulating different processes featuring the berry ripening progression.

NAC61 is highly expressed during post-harvest berry dehydration and its expression pattern is closely related to sugar concentration. The ectopic expression of *NAC61* in *Nicotiana benthamiana* leaves determines low stomatal conductance, high leaf temperature, tissue collapse and a higher relative water content. Transcriptome analysis of grapevine leaves transiently overexpressing *NAC61,* and DNA affinity purification and sequencing analyses allowed us to narrow down a list of NAC61-regulated genes. Direct regulation of the stilbene synthase regulator *MYB14*, the osmotic stress-related gene *DHN1b*, the *Botrytis cinerea* susceptibility gene *WRKY52* and the *NAC61* itself, is validated. We also demonstrate that NAC61 interacts with NAC60, a proposed master regulator of grapevine organ maturation, in the activation of *MYB14* and *NAC61* expression. Overall, our findings establish NAC61 as a key player in a regulative network that governs stilbenoid metabolism and osmotic, oxidative and biotic stress responses in grape berry during late-and post-ripening.

**Highlights:** NAC61 regulates stilbene biosynthesis and abiotic/biotic stress responses that hallmark late-and post-ripening developmental stages in grapevine berry. NAC61 participates in a NAC60-dependent regulatory network, also triggering its self-activation.

## INTRODUCTION

Fruit ripening is an irreversible highly regulated process involving physiological and biochemical changes maximizing fruit organoleptic traits to attract herbivores and facilitate seed dispersal (Giovannoni, 2004). The main changes that take place during ripening include fruit de-greening, colored pigment accumulation, textural changes (leading to softening) and composition changes, such as the depletion of organic acids and the accumulation of sugars and aroma compounds. This complex program peaks when mature seeds are ready to be dispersed. Nonetheless, at advanced ripening, tissue softening, and eventual decay make fruits susceptible to attack by opportunistic pathogens and, consequently, an enhancement of the constitutive defense against pathogens is inherent in the ripening program. Several processes taking place during ripening, such as chloroplast and cell wall disassembly, reactive oxygen species (ROS) increase, protein degradation, and the activation of the secondary metabolism, resemble senescence-like processes (Gómez *et al*., 2014). However, some specific metabolic activities and the fact that only subsets of senescence-related genes are activated during ripening, put forward that, albeit partially recruiting processes and metabolisms typically associated to senescing tissues, ripening is a distinct process that precedes and may predispose the fruit to subsequent senescence (Forlani *et al*., 2019; Gapper *et al*., 2013).

To reach its final composition, the grapevine berry, a typical non-climacteric fruit, undergoes a developmental process comprising a vegetative and a ripening growth phase (Zenoni *et al*., 2021). The vegetative phase involves pericarp growth due to rapid cell division and the accumulation of organic acids, tannins and other phenolic compounds. The ripening phase features several physical, physiological and compositional changes such as pericarp tissue softening, cell expansion, loss of organic acids, anthocyanin accumulation in the skin, and the progressive accumulation of sugars, reaching levels normally over 20% in the juice (Conde, 2007). Several studies revealed that the onset of ripening, known as veraison, coincides with a profound transcriptomic rearrangement featuring the rapid downregulation of genes strongly expressed during the vegetative phase of berry development and the upregulation of genes participating in the ripening program (Fasoli *et al*., 2012; Fasoli *et al*., 2018; Massonnet *et al*., 2017). Moreover, additional extensive transcriptomic and metabolic changes have been shown to occur in berries of clusters left on the vine beyond ripening or harvested and placed in dehydrating rooms (Zamboni *et al*., 2008; Zenoni *et al*., 2016), thus revealing that the developmental program of the grape berry is not terminated at the fruit commercial ripening. These studies identified transcription factors (TFs) as putative master regulators of the grape berry developmental progression and the metabolisms featured in each developmental stage (Fasoli *et al*., 2018; Massonnet *et al*., 2017; Palumbo *et al*., 2014; Zenoni *et al*., 2016). Among these, several members of the NAC (NAM/ATAF1/CUC2) TF gene family are included. NACs represent plant-specific TFs with a wide range of activities during plant and fruit development (Forlani *et al*., 2021; Olsen *et al*., 2005). The tomato non-ripening (NOR) is the first NAC TF described as a master regulator of fruit ripening (Gao *et al*., 2020; Kumar *et al*., 2018; White, 2002), recently shown to play a role also in leaf senescence (Ma *et al*., 2019). SlNAC1 (also known as

SlNAC033) has been shown to have a role in heat and chilling tolerance (Liang *et al*., 2015; Ma *et al*., 2013), in the defense against bacterial pathogens (Huang *et al*., 2013) and in fruit softening and pigmentation (Ma *et al*., 2014; Meng *et al*., 2016). The Arabidopsis AtNAP (NAC-like, Activated by AP3/PI, ANAC029), has been shown to promote both silique maturation and leaf senescence (Guo and Gan, 2006; Kou *et al*., 2012). The strawberry FaNAC035 was demonstrated to regulate ripening by controlling fruit softening as well as pigment and sugar accumulation, through the regulation of abscisic acid (ABA) biosynthesis and signalling, and cell wall degradation and modification (Martín-Pizarro *et al*., 2021). The NAC TFs are also involved in drought and oxidative stress responses (Balazadeh *et al*., 2011; Mao *et al*., 2012; Puranik *et al*., 2012; Tran *et al*., 2004) and in the regulation of ROS metabolism (Fang *et al*., 2015).

In grapevine, *NAC33* and *NAC60* have been functionally investigated as putative master regulators of the vegetative-to-mature transition in several plant organs (D’Inca *et al*., 2021; D’Inca *et al*., 2023). *NAC33* plays a major role in leaf and fruit, terminating photosynthetic activity and organ growth, while for *NAC60*, able to complement the *nor* mutant phenotype in tomato, a dual role as an orchestrator of both ripening-and senescence-related processes has been proposed. Moreover, it has been shown that NAC60 homodimer is the prevalent form in berries during ripening, although its ability to form heterodimers with NAC03 and NAC33 has also been demonstrated, suggesting the existence of a NAC60-NAC regulatory network (D’Inca *et al*., 2023). Interestingly, *NAC60*, together with the still uncharacterized *NAC61*, were also identified as markers of post-harvest dehydration (Zenoni *et al*., 2016).

Here, we report the functional characterization of the grapevine NAC TF *NAC61*, providing evidence of its role in the regulation of berry late-and post-ripening processes. We studied the *NAC61* function through the study of its expression and co-expression pattern, its ectopic overexpression in *Nicotiana benthamiana*, its transient overexpression in grapevine plants, and by performing DNA Affinity Purification and sequencing (DAP-seq). We identified direct targets of NAC61 and demonstrated its ability to activate genes acting in stilbene biosynthesis and osmotic, heat and oxidative/biotic stress responses. We also investigated the *NAC61* upstream regulation, demonstrating its own and NAC60-mediated activation, showing that abiotic and biotic factors may influence its expression.

## MATERIAL AND METHODS

### Plant material

*Vitis vinifera* cv. ‘Thompson Seedless’ plantlets and embryogenic *calli*, and *Nicotiana benthamiana* plants were grown as previously described (Amato *et al*., 2019). *Vitis vinifera* cv. ‘Syrah’ fruiting cuttings were propagated as previously described (Mullins and Rajasekaran, 1981).

### Isolation and cloning

The *NAC61* coding sequence (CDS) was isolated from cv. ‘Syrah’ ripening berry cDNA, and the regulative region of *NAC61*, *DHN1b*, *MYB14* and *WRKY52* were isolated from *Vitis vinifera* cv. ‘Syrah’ genomic DNA. cDNA and genomic DNA were extracted from *Vitis vinifera* cv. ‘Syrah’ fruiting cuttings prepared as previously described (D’Inca *et al*., 2021). Amplification was performed by using the KAPA HiFi DNA polymerase (KAPA Biosystems, Wilmington, MA, USA) and primers sets listed in **Supplementary Table S1.** The isolated sequences were directionally cloned into the *pENTR/D-TOPO* Gateway entry vector (Invitrogen, Waltham, MA, USA) and transferred by site-specific LR recombination into a specific binary vector (http://www.vib.be/en/research/services/Pages/Gateway-Services.aspx). For *N. benthamiana* and *Vitis vinifera* cv. ‘Thompson Seedless’ plantlets agroinfiltration, the *NAC61* CDS was transferred into the *pK7GW2.0* binary overexpression vector. For the dual luciferase reporter assay (DLRA), the *NAC61*, *DHN1b*, *MYB14* and *WRKY52* target gene regulative regions were transferred into the *pPGWL7.0* reporter vector to control the *Firefly* luciferase (*LUC*) expression. For the bimolecular fluorescence complementation assay (BiFC) assay, the *NAC61* sequence was transferred into the *pnYGW* vector.

The *NAC60* CDS isolation, cloning into the *pENTR/D-TOPO* entry vector and the site-specific LR recombination into the *pK7GW2.0* binary overexpression vector was previously performed (D’Incà *et al*. 2023).

### Transient overexpression

The *pK7WG2.0* vectors containing *35S:NAC61* or a noncoding sequence (control, CTRL) were transferred to *Agrobacterium tumefaciens* strain C58C1 by electroporation. For transient expression in grapevine cv. ‘Thompson Seedless’, five-week-old *in vitro* plantlets were vacuum infiltrated as previously described (Amato *et al*., 2016) and molecular analyses were carried out on leaf samples collected seven days after the agroinfiltration. For transient expression in *N. benthamiana*, fully expanded leaves were syringe infiltrated and phenotypic analysis was carried out over three days after agroinfiltration.

### Stomatal conductance and thermal camera measurements

The measurements were performed on five-weeks-old healthy *N. benthamiana* plant overexpressing *35S:NAC61* and the control. The stomatal conductance measurements were carried out for six days post infiltration by using a portable leaf porometer (SC-1, METER Group, Inc., Pullman, WA, USA). Three biological replicates (different plants), each with three technical replicates (different leaves), were performed for each sample, bringing to nine replicates. The thermal images were taken for six days post infiltration with a thermal camera (FLIR E6 Wifi, FLIR Systems, Sweden). Six biological replicates (different plants), each with three technical replicates (different leaves), were performed for each sample, bringing to 18 replicates.

### Relative water content (RWC) measurement

The experiment was performed on five-weeks-old healthy *N. benthamiana* plant overexpressing *35S:NAC61* and the control as previously described (Xu *et al*., 2022). Three biological replicates (different plants), each with three technical replicates (different leaves), were performed for each sample, bringing to nine replicates. For measurement of RWC, leaves from each plant were sampled two days after agroinfiltration. Leaves were immediately weighted and recorded as ‘fw’, then leaves were immersed in distilled water for two hours at room temperature. The weight of hydrated leaves was measured and recorded as ‘w’; the dry weight was measured after 24 hours of drying at 60°C and indicated as ‘dw’. The RWC was measured as [(fw-dw)/(w-dw)]*100. The evaluation of local water accumulation was performed by the agroinfiltration of *35S:NAC61* and control vectors in delimited portion of the same leaf. Three biological replicates (different plants), each with three technical replicates (different leaves), were performed, bringing to nine replicates for each sample. Infiltrated leaves were daily inspected and pictures of the abaxial face were taken to observe the progression of the phenotype within agroinfiltrated tissues.

### Ion leakage

Ion leakage was performed according to Imanifard *et al*., (2018) on five-weeks-old healthy *N. benthamiana* plant overexpressing *35S:NAC61* and the control. Six biological replicates (different plants), each with three technical replicates (different leaves), were performed for each sample, bringing to 18 replicates. The leaf discs (∼5 mm in diameter) were collected after two days from the infiltration and immersed in 50 mL of non-ionic double-distilled water for 30 minutes at 90 rpm and 25°C to eliminate ions released because of physical damage. The 18 leaf discs from each sample were distributed into three wells of a multi-well plate containing 2 mL of distilled water per well (one replicate per well, the six discs are each from an independent plant to avoid plant specific effects). The conductivity of the solution was measured using a conductivity meter (Horiba scientific, Edison, NJ, USA) immediately after the plate preparation and during 24 hours of shaking at 90 rpm under constant light (50 μmol/m2 s) at 25 °C.

### 3,3’-diaminobenzidine (DAB) assay

The staining was performed on five-weeks-old healthy *N. benthamiana* plant overexpressing *35S:NAC61* and the control as previously described (Daudi and O’Brien, 2012). Three biological replicates (different plants), each with three technical replicates (different leaves), were performed for each sample, bringing to nine replicates. The leaf discs (∼2 cm in diameter) were collected and used for the assay. A DAB solution was used to evaluate the production of H_2_O_2_ after two days from the agroinfiltration. The assay was performed in 12-well plates and the DAB staining solution was vacuum infiltrated. The plate was covered with aluminum foil and incubated for five hours at 100 rpm shaking at room temperature. At the end of the staining and to degrade all the chlorophylls, the disks were placed in falcon tubes containing 25 mL of ethanol 80%. Finally, the staining quantification was analyzed through ImageJ software (https://imagej.net/ij/index.html).

### Real Time quantitative Polymerase Chain Reaction (RT-qPCR)

For gene expression analysis RNA was isolated from 100 mg of leaf ground material (for the cv. ‘Thompson Seedless’ transcriptomic analysis) and 200 mg of berry ground material (for the expression analysis on cv. ‘Müller-Thurgau’ drying berries), using the Spectrum Plant Total RNA kit (Merck KGaA, Darmstadt, Germany). Gene expression was determined by RT-qPCR as previously described (Zenoni *et al*., 2011) using the primer sets listed in **Supplementary Table S1.** Each value corresponds to the mean ± SD of three replicates relative to the *UBIQUITIN1* (*VIT_16s0098g01190*) internal control. *Vitis vinifera* cv. ‘Corvina’ mature and low/high temperature dried berries cDNA was previously prepared by (Shmuleviz *et al*., 2023).

### Transcriptomic analysis on *NAC61* transient overexpressing grapevine plants

The microarray analysis was performed with the RNA used for RT-qPCR. For transient expression, the three most highly overexpressing plants and three control lines were selected and used as biological replicates. The cDNA synthesis, labelling, hybridization and washing were performed according to the Agilent Microarray-Based Gene Expression Analysis Guide (v.6.5). Each sample was hybridized to an Agilent custom microarray four-pack 44K format (Agilent Sure Print HD 4X44K 60-mer; cat. no. G2514F-048771) (Dal Santo *et al*., 2016) and scanned using an Agilent Scanner (G2565CA; Agilent Technologies, Santa Clara, CA, USA). Feature extraction and statistical analysis of the microarray data was conducted by using the Limma package in R (Ritchie *et al*., 2015). p-values were normalized using the Benjamini–Hochberg correction (Benjamini and Hochberg, 1995) and the differentially expressed genes (DEGs) were identified by adjusted p value <0.1 and selected by fold change (FC) >|1,5|.

### DAP-seq

Genomic DNA was extracted from 1 g of ground young cv. ‘Syrah’ leaves and the Illumina libraries were prepared as previously described (Galli *et al*., 2018). The *NAC61* sequence was transferred from the *pENTR/D-TOPO* to the Gateway-compatible destination vector *pIX-HALO* (Bartlett *et al*., 2017). The HALO-NAC61 and GST-HALO (used as negative control) fusion proteins were translated *in vitro* using the TNT^R^ SP6 coupled reticulocyte lysate system (Promega). Two replicates were used for the TF and input libraries. The DAP-seq was performed according to previously described procedure (Galli *et al*., 2018) and a total of 7.0 (control replicate 1), 5.6 (control replicate 2), 24 (NAC61 replicate 1) and 22 (NAC61 replicate 2) millions reads were obtained. DAP-seq bioinformatic analysis was performed as previously described (D’Inca *et al*., 2023; Orduna *et al*., 2022; Orduña-Rubio *et al*., 2023). Briefly, DAP-seq libraries were aligned to the PN40024 12X.v2 reference genome using bowtie2 (Langmead and Salzberg, 2012), with post-processing to remove reads with a MAPQ score lower than 30. Peak detection was performed using GEM peak caller (Guo *et al*., 2012) version 3.4 with the 12X.v2 genome assembly using the following parameters: ‘–q 1 –t 1 –k_min 6 –kmax 20 –k seqs 600 – k_neg_dinu_shuffle’, limited to nuclear chromosomes. The replicates were analyzed as multi-replicates with the GEM replicate mode. Detected peaks were associated to closest gene model from the PN40024 v1 on the 12X.0 assembly transposed to the 12X.2 assembly annotation file using the BioConductor package ChIPpeakAnno (Zhu *et al*., 2010a) with default parameters.

### DLRA

The *pK7WG2.0* vectors containing the *NAC61* and *NAC60* CDS and the *pPGWL7.0* vectors harboring the *DHN1b*, *MYB14* and *WRKY52* regulative regions were transferred to *Agrobacterium tumefaciens* C58C1 strain by electroporation. The DLRA was performed on three fully expanded infiltrated *N. benthamiana* leaves from three different plants, as previously described (Cavallini *et al*., 2015). The assay was performed on fresh leaf disks collected 72 hours after Agrobacterium-mediated infection and following the manufacturers instruction (Promega). A reference vector overexpressing the *Renilla reniform*is luciferase (*REN*) was used to normalize LUC luminescence. REN and LUC luminescence were detected using a Tecan Infinite ® M200 PLEX instrument. Each test was performed in biological triplicate and each value was measured in triplicate.

### Grapevine protoplast transfection and BiFC assay

For the BiFC assay, the *NAC61* and *NAC60* CDS were cloned into the *pnYGW* and *pGWcY* Gateway vectors, respectively. *V. vinifera* cv. ‘Thompson Seedless’ protoplasts were isolated from embryogenic calli and transfected (Bertini *et al*., 2019), cultured into multi-well plates in the dark at 25°C and analyzed one day after transfection. The yellow fluorescent protein (YFP) signal was detected using a Leica TCS SP5 AOBS confocal microscope (Argon laser, 514 nm excitation source, 550-570 nm collection bandwidth, auto gain).

### *Vitis vinifera* cv. ‘Müller-Thurgau’ berry post-harvest dehydration and noble rot induction

About 405 kg of berry bunches of *V. vinifera* cv. ‘Müller-Thurgau’ were harvested in August 2017 in Custoza (Italy) when the soluble solids content was 18.25 ± 0.05° Brix and transferred in a ventilated dehydration facility of the farm ‘La Prendina’ in Monzambano (Mantova, Italy) for dehydration under controlled conditions (14-15°C, 53–60% relative humidity) reached by DEUM 5 HP machine (Sordato, Verona, Italy). Three randomly selected replicate berries were regularly sampled to determine the soluble solids content using a DBR35 digital refractometer (Giorgio Bormac, Carpi, Italy). Moreover, three dedicated *plateaux* were weighed regularly using a CH50K50 electronic balance (Kern, Balingen, Germany). After 29 days of dehydration, half of the boxes were covered with plastic film and water-filled trays were placed inside to increase the relative humidity and induce noble rot (Negri *et al*., 2017), while the remaining plastic boxes (control berries) were left under normal dehydrating conditions. The two different environmental conditions were imposed for an additional 28 days. The relative humidity (RH), in both conditions, were monitored using Hobo Pro v2 sensors connected to data loggers (Onset Computer Corporation, Bourne, MA, USA), and berry pH were weekly measured on three biological replicates of 50 berries each. Control and noble rot-induced berries were sampled in biological triplicate for transcriptomic analysis 7, 21 and 28 days (t1, t2 and t3, respectively) from the induction. For the glycerol to d-gluconic acid ratio measurements, a glycerol assay kit (Merck), and a d-gluconic acid/d-glucono-δ-lactone assay kit (Megazyme International), were used according to the manufacturer’s instructions 1 g of berry pericarp powdered material was diluted in 10 mL of buffer containing 500 µL Carrez 1 solution, 500 µL Carrez 2 solution, and 1 mL 0.1 m NaOH topped up to 10 mL with water and filtered with standard filter paper. After a treatment with polyvinylpolypyrrolidone to remove colored solutes, samples have been additionally filtered.

### Gene Co-expression Networks (GCNs), binding motif comparison and promoter analyses

*NAC61* GCNs were extracted from the AggGCNs app (Orduña-Rubio *et al*., 2023). Gene set enrichment analysis conducted in this study was conducted with the gprofiler2 R package (Kolberg *et al*., 2020), using the MapMan manually curated annotation described (Orduña-Rubio *et al*., 2023) with default settings. A significance threshold of 0.05 was chosen for p-values adjusted with the Benjamini–Hochberg correction procedure (Benjamini and Hochberg, 1995). Enriched sequences found in NAC61 binding sites were compared with those found in *Arabidopsis thaliana* using the RSAT Plants NGS-ChIP-seq Peak-Discovery software (https://rsat.eead.csic.es/plants/peak-motif_form.cgi) (Santana-Garcia *et al*., 2022). The top 600 best scored peak sequences (-50bp<Peak center<+50bp) were retrieved from the DAP-seq analysis and used with default parameters. Most significant, frequent and middle-centered motifs were selected. An untargeted binding discovery analysis and the targeted binding comparison analysis was also performed on the *NAC61* promoter using the RSAT Plants Motif discovery oligo-analysis (words) software (https://rsat.eead.csic.es/plants/oligo-analysis_form.cgi), using the default parameters (selecting ‘*Vitis vinifera* PN40024.v4.55’ as organism), and the RSAT Plants Pattern Matching Matrix-scan (https://rsat.eead.csic.es/plants/matrix-scan_form.cgi), for surveying the ‘Arabidopsis PBM’, ‘Athamap’ and ‘Cistrome’ dataset.

## RESULTS

### *NAC61* is upregulated in post-veraison stages and correlates with osmotic stress in grape berries

The *NAC61* expression pattern, according to the global gene expression atlas of *Vitis vinifera* (Fasoli *et al*., 2012), shows an increase during the berry development and, additionally, in other organs such as seeds, rachis, stems and roots, while a weak expression is found in flower organs (**Fig. 1A**). In berry tissues, the *NAC61* expression level sharply increases at veraison and shows a second step of up-regulation after harvest, reaching the highest expression level at the end of the post-harvest dehydration process (post-harvest withering stages; **Fig. 1A**; **Supplementary Fig. S1A-B**). By examining a transcriptomic dataset from a Genotype x Environment (GxE; Dal Santo *et al*., 2018) study, we observe that *NAC61* belongs to a stage-specific cluster of genes, whose expression increases after veraison and is thus poorly affected by environmental conditions (**Supplementary Fig. S1C**). We then investigated the relation between the *NAC61* expression level and the sugar content in berry, finding a high positive correlation occurring during ripening when the berry exceeds the sugar level of 15-18 °Brix (**Fig. 1B**). Such close relation between sugar concentration and *NAC61* expression is maintained until the end of ripening and is also observed in berries during post-harvest withering (PHW), in which the highest expression level is coincident with the highest sugar concentration, independently of the genotype considered (**Fig. 1C**). The inspection of the transcriptomic dataset from a study, aiming specifically at dissecting the effect of time and dehydration level in post-harvest dehydrating berries (Zenoni *et al*., 2020), reveal that *NAC61* expression is more correlated with the grape weight loss level (an indirect measure of sugar concentration) than time, further strengthening the above-reported observations (**Fig. 1D**).

**Figure 1.**
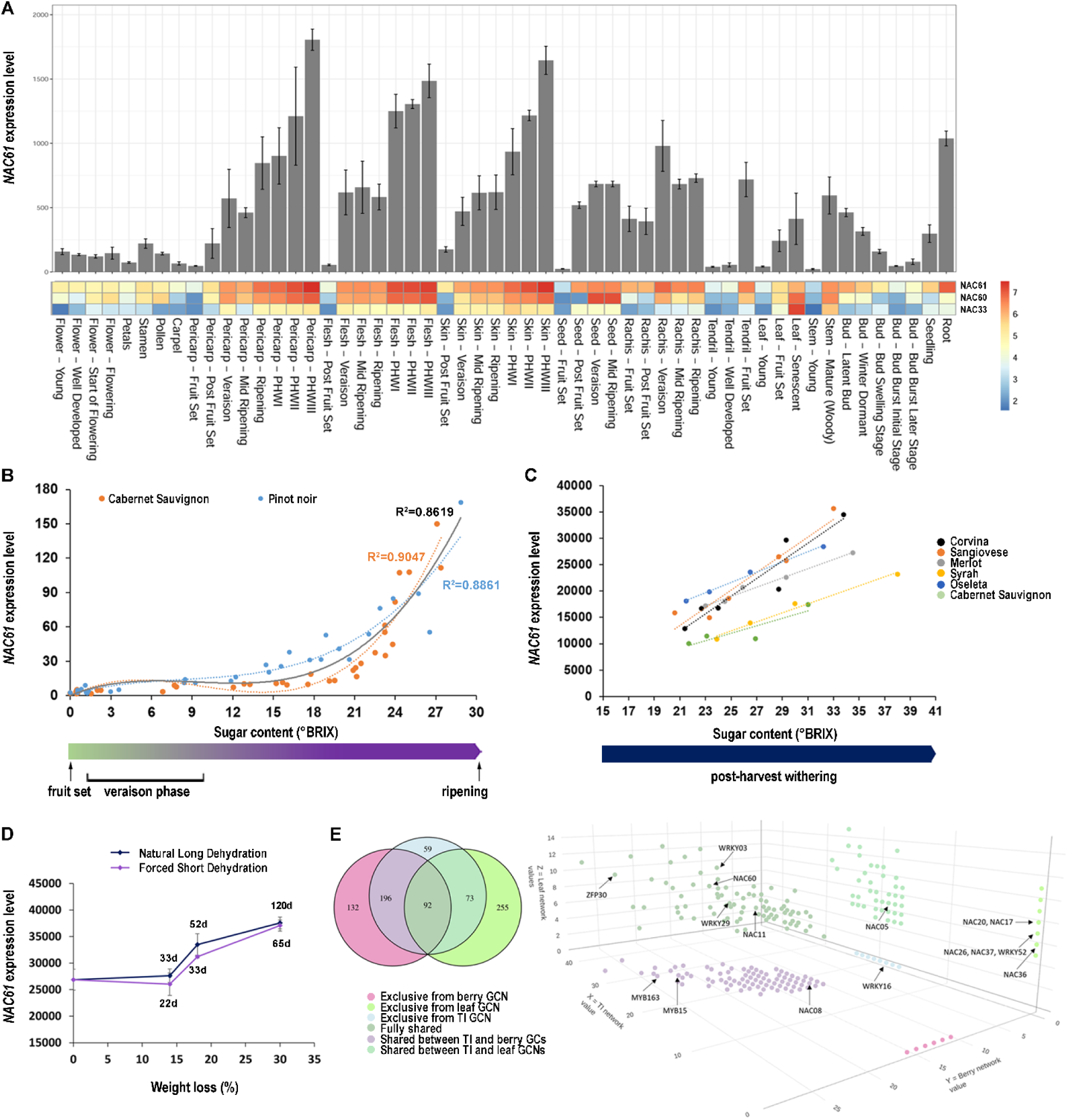
*NAC61* expression analysis. (A) *NAC61* expression behavior in grapevine organs throughout development (bar plot) compared in the heatmap (logaritmic value) with that of *NAC60* and *NAC33.* The data were retrieved from the atlas transcriptomic dataset of cv. ‘Corvina’ (Fasoli *et al*., 2012) and each value represents the mean (± SD) of three biological replicates. (B) Correlation between *NAC61* expression level and sugar content in grape berries sampled from fruit set to maturity in cv. ‘Cabernet Sauvignon’ and cv. ‘Pinot noir’ (Fasoli *et al*., 2018). Black line represents the trend of the averaged values of the two varieties. (C) Correlation between *NAC61* expression level and sugar content in grape berries sampled during post-harvest dehydration in six different varieties (Zenoni *et al*., 2016). (D) Correlation between *NAC61* expression level and berries weight loss in cv. ‘Corvina’ berries sampled during traditional long and forced short post-harvest dehydration processes (Zenoni *et al*., 2020). Expression values were determined by microarray and each value represents the average from three biological replicates (± SD). (E) *NAC61* GCNs based on berry, leaf and TI datasets. To the left the Venn diagram showing exclusive and shared genes by the three datasets and to the right the 3D plot of co-expressed genes in which NAC, WRKY and ZIP members already described for their involvement in berry ripening and/or stress responses, are highlighted.

Extracting the *NAC61* gene-centered networks from berry (67 experiments), leaf (42 experiments) and tissue-independent (TI; 131 experiments) datasets through the AggGCN app within the VitViz platform (http://www.vitviz.tomsbiolab.com/), we identify a total of 810 *NAC61* co-expressed genes mainly belonging to ‘Transcription regulation’ and ‘Transcription factors’ functional categories (**Supplementary Dataset S1; Supplementary Fig. S2**). Most of the NAC61 co-expressed TFs belong to NAC, zinc finger, WRKY and MYB families (**Fig. 1E; Supplementary Dataset S1)**. Moreover, we find 98 genes previously defined as key regulators of berry ripening and 575 genes, 71% of the total, which are differentially modulated during the post-harvest dehydration process (Fasoli *et al*., 2018; Massonnet *et al*., 2017; Palumbo *et al*., 2014; Zenoni *et al*., 2016; **Supplementary Dataset S1**). Accordingly, the stilbene synthases (STSs) regulators *MYB15, WRKY03* and *WRKY43* are also co-expressed with *NAC61*.

The NAC family multispecies phylogenetic tree (https://tomsbiolab.com/wp-content/uploads/2021/10/Fig.-S4.png; **Supplementary Fig. S3**) reveal that *NAC61* is located close to *ANAC046* and *NAC33*, both involved in the senescence process (D’Inca *et al*., 2021; Oda-Yamamizo *et al*., 2016), to *ORS1*, coding for an H2O2-responsive NAC TF also controlling senescence in Arabidopsis (Balazadeh *et al*., 2011), and to *OsNAC2*, a positive regulator of drought and salt tolerance through ABA-mediated pathways in rice (Jiang *et al*., 2019).

### *NAC61* heterologous expression induces water-soaking-like phenotype and programmed cell death in *Nicotiana benthamiana* leaves

The role of *NAC61* is studied through transient heterologous expression in *N. benthamiana* plants. Three days after agroinfiltration, transgenic leaves show leaf tissue collapse with loss of turgor (**Fig. 2A**). To highlight the effect of NAC61, leaves were simultaneously agroinfiltrated with *NAC61,* the control vector and the agroinfiltration buffer. At two days post infiltration (48 h) part of the leaves near the site of *NAC61* infiltration show a darker color, suggesting a local accumulation of water that precedes tissue collapse observed at three (72 h) and four (96 h) days after infiltration (**Fig. 2A)**.

**Figure 2.**
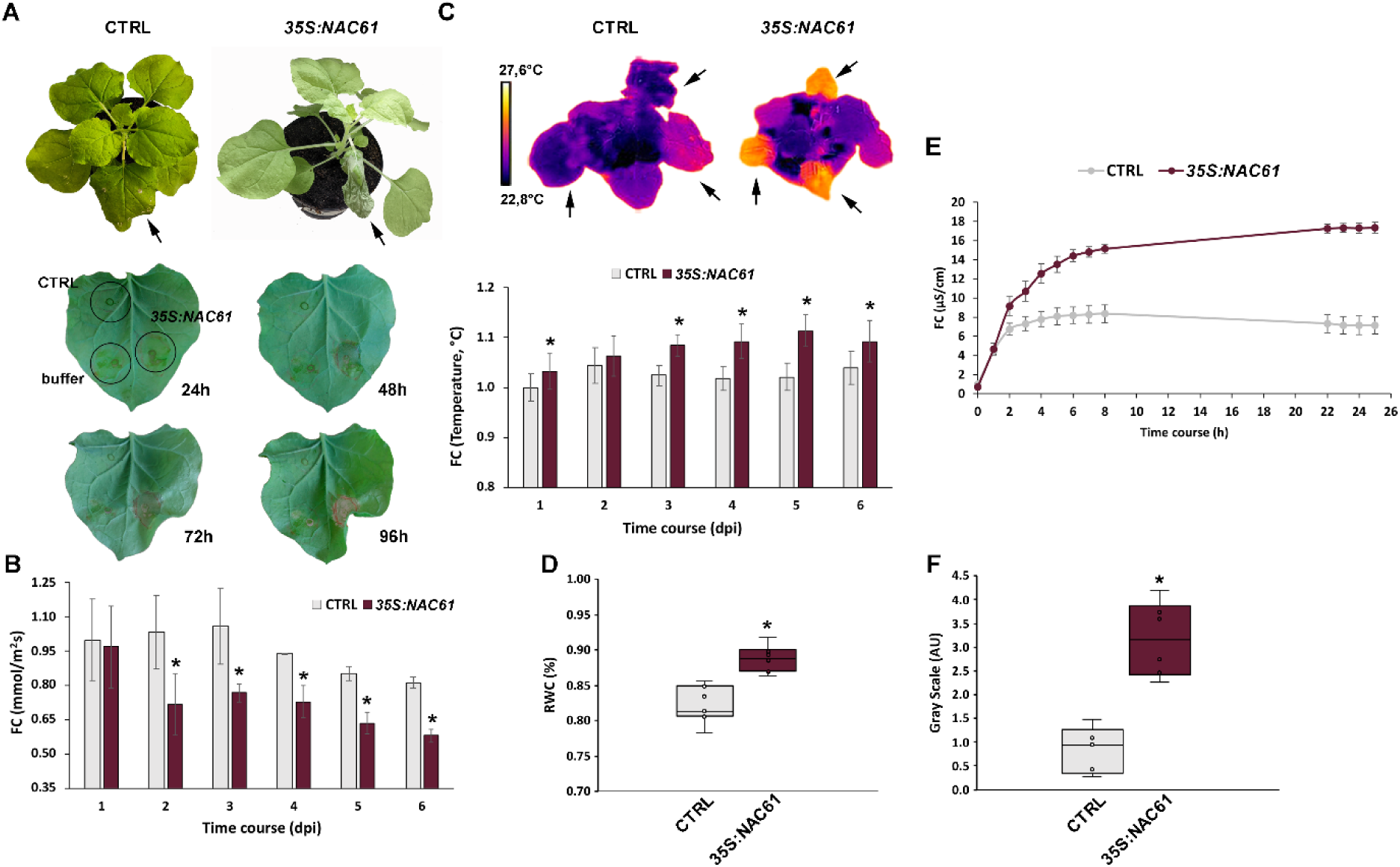
*NAC61* ectopic expression in *Nicotiana benthamiana* plants. (A) Control and *NAC61*-expressing *Nicotiana benthamiana* plant (on the top) three days after infection, and spot-infiltrated leaves after 24, 48, 72 and 96 hours. (B) Stomatal conductance measurements in the *NAC61*-expressing leaves compared to the control. (C) Thermal camera visualization and measurements in the *NAC61*-expressing leaves compared to the control. (D) RWC measurements in *NAC61*-expressing leaves compared to the control, two days after agroinfiltration (E) Ion leakage measurements in *NAC61*-expressing leaves compared to the control from 24 to 48 hours after agroinfiltration. (F) DAB staining determining H_2_O_2_ accumulation in *NAC61* expressing leaves compared to the control, two days after agroinfiltration. Each value represents the mean (± SD) of three biological replicates tested in technical replicate (*n=* 3). Asterisks indicate statistically significant differences (*, p < 0.01; *t-*test).

Interestingly starting from two days after *NAC61*-agroinfiltration, transgenic leaves display a lower stomatal conductance compared to the control (**Fig. 2B**). Consistently, a significantly higher leaf temperature was registered (**Fig. 2C).** To investigate if *NAC61* expression favor water retention within plant tissue we measured the RWC in transgenic and control leaves at two days after infiltration. The analysis revealed a significant increase of water content in NAC61-overexpressing leaves (**Fig. 2D)** resembling a water-soaking phenotype.

Moreover, in line with the cell death observed at whole leaf level at four days post infection (Fig. 2A), *NAC61*-expression significantly increases ion leakage already after two days, suggesting a possible loss of membrane integrity (**Fig. 2F**). Finally, a higher level of H_2_O_2_ is observed in the *NAC61*-expressing plants in comparison to the control (**Fig. 2G**), in line with the above-described phenotype, as H_2_O_2_ is a key ROS involved in both stomatal closure and programmed cell death (Liu and Zhang, 2021).

### Transient *NAC61* overexpression in *Vitis vinifera* affects stilbenoid-related gene expression

Transiently *NAC61*-overexpressing grapevine cv. ‘Thompson Seedless’ plants (**Supplementary Fig. S4A**) display 1157 DEGs compared to the control plants (**Supplementary Dataset S2**). Among DEGs, 530 genes are upregulated and 627 are downregulated. No phenotypic alterations are observed in *NAC61*-overexpressing leaves. Gene category MapMan distribution and DEGs enrichment analysis reveal that upregulated genes are mainly involved in ‘Secondary metabolism’, in particular ‘Stilbenoid biosynthesis’, and in ‘Oxidoreductases activity’, in particular ‘Laccases’, while downregulated genes are mainly represented by ‘Photosynthesis-related mechanisms’ and other primary metabolism-related processes, such as ‘Lipid and Carbohydrate metabolisms’ (**Fig. 3A**; **Supplementary Dataset S3)**.

**Figure 3:**
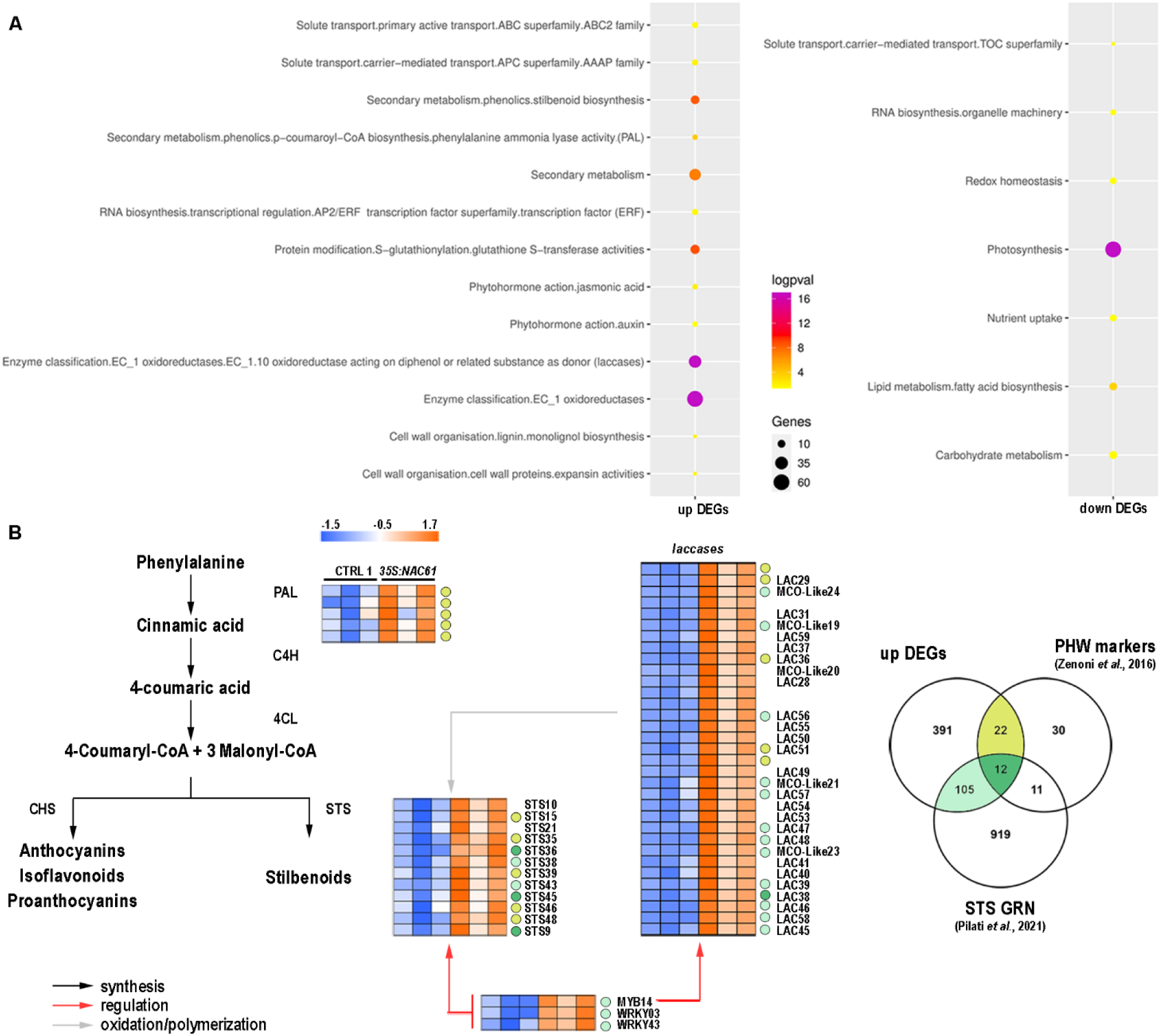
Transcriptomic responses to *NAC61* overexpression in the cv. ‘Thompson seedless’ grapevine leaves. (A) Functional enrichment analysis of upregulated and downregulated DEGs. (B) Heatmap of upregulated DEGs involved in phenylpropanoid synthesis, regulation and modification (**Supplementary Dataset S2**). Markers of the PHW process (Zenoni *et al*., 2016) and genes belonging to the STS GRN (Pilati *et al*., 2021) are highlighted according to the Venn diagram color code.

We highlight that NAC61 highly affects the expression of five *phenylalanine ammonia lyases* (*PALs*), corresponding to the first and committed step in the phenylpropanoid pathway, and 12 *STSs*, the key enzymes leading to stilbenoid biosynthesis (**Fig. 3B**). The five *PALs* and eight of the 12 *STSs* were previously described as markers of post-harvest dehydration (Zenoni *et al*., 2016). Upregulated DEGs also include 33 *laccases* (*LACs*), proposed to be involved in the oxidative polymerization of phenolic compounds (Keylor *et al*., 2015), and six of which were also described as markers of post-harvest dehydration (**Fig. 3B**). Overall, 34 out of the 75 molecular markers of the post-harvest berry dehydration are found upregulated by NAC61.

By inspecting the recently proposed *STS* Gene Regulatory Network (GRN), built by OneGenE tool (Pilati *et al*., 2021), we find 117 of the 530 upregulated genes (**Fig. 3B)**, including the stilbenoid regulators *MYB14, WRKY03* and *WRKY43*. Moreover, 13 out of the total upregulated *LACs* are also found in the *STS* GRN, ascribing their potential role in stilbene polymerization (i.e. production of viniferins), as previously suggested (Orduna *et al*., 2022; Pilati *et al*., 2021; Zenoni *et al*., 2016). The upregulation of *MYB14* and a *laccase* gene (*LAC25*) in the *NAC61-*overexpressing plants is validated by RT-qPCR (**Supplementary Fig. S4B)**. Of note, the ectopic expression of *NAC61* induces also *NAC61* itself.

### Examination of NAC61 cistrome for identifying NAC61 high confident targets (HCTs)

To identify putative direct targets of NAC61, we inspect its genome-wide binding landscape (cistrome) by carrying out DAP-seq. We identify 8558 binding events assigned to 6734 genes (**Fig. 4A**). The distribution of NAC61 DNA-binding events, with respect to their position from the transcription start site (TSS) of the identified genes, shows the preferential localization in proximal upstream regions and inside genes (**Fig. 4A**; **Supplementary Dataset S4**). Within the ‘inside gene’ category, most binding events are found at the very start of the gene feature, i.e. close to 100 pb. By inspecting the 600 top scoring peaks, we identify the major binding motif [CA(C/A)G(C/T)(A/C)A] (**Fig. 4B**), correlated with *A. thaliana* ANAC46, controlling cell death during leaf senescence (Huysmans *et al*., 2018), ANAC55 involved in ABA and jasmonic acid responses (Jiang *et al*., 2009), and ANAC047, the NAC60 closest homologue (D’Inca *et al*., 2023; **Supplementary Fig. S5**) according with RSAT Plants phylogenetic footprints. To identify the putative targets of NAC61, we focus on genes for which the TF bound to their promoter region (from -3kb to +100bp to the TSS), thus identifying 2471 peaks and 2263 unique genes (**Supplementary Dataset S4**). The use of the MapMan ontology of these genes highlights a functional enrichment in ‘Transcriptional regulation (*zinc fingers*, *NACs*, *MYBs*, *ERFs*)’, ‘Solute transport’, ‘Protein homeostasis’ and ‘External stimuli and pathogen response’ descriptors (**Fig. 4C**).

**Figure 4:**
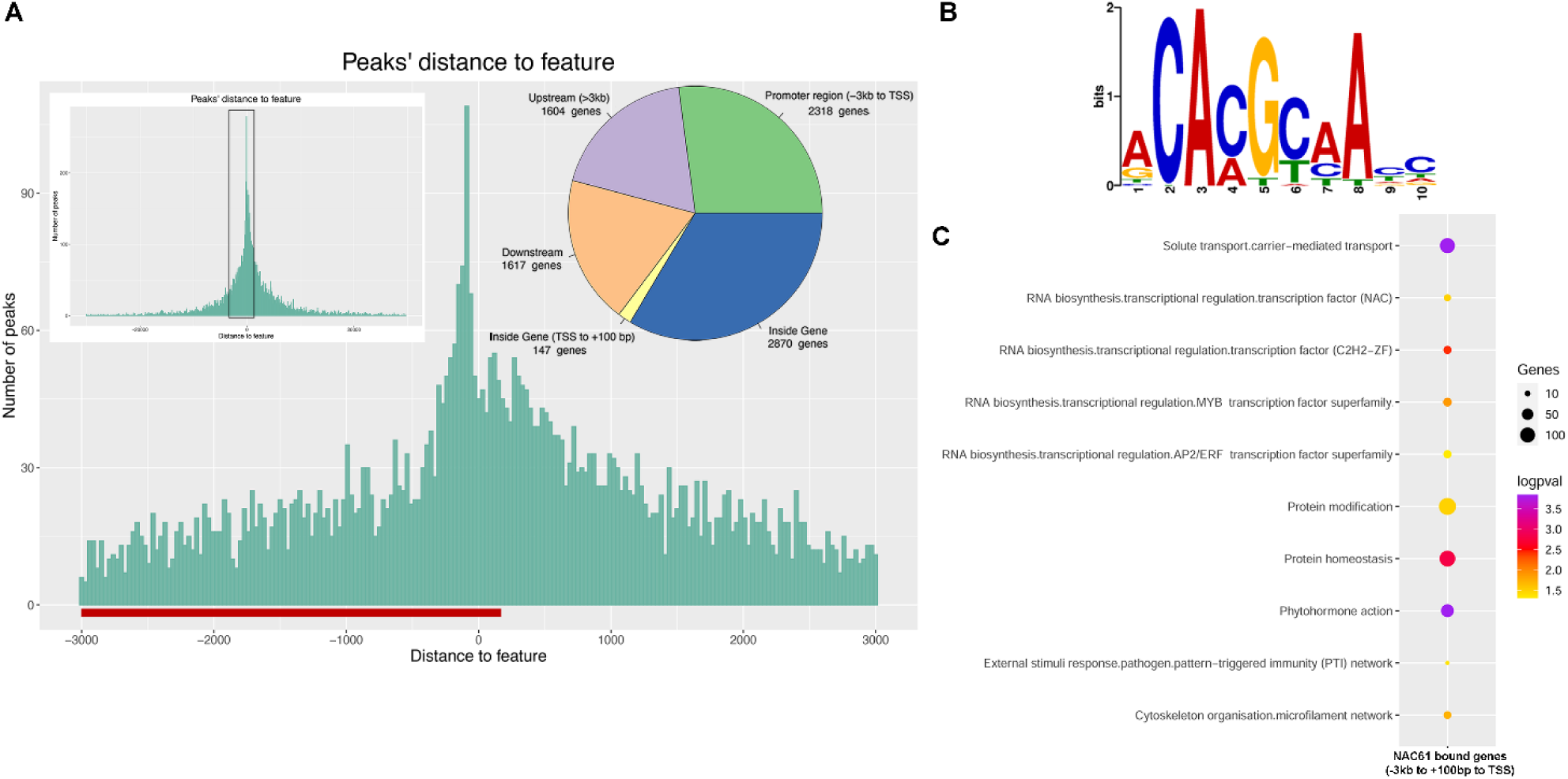
NAC61 DAP-seq analyses. (A) Distribution of NAC61 DNA binding events with respect to their position from the TSS of their assigned genes. Distribution of peak positions is represented with a pie chart. (B) *De novo* forward binding motif obtained from the inspection of the top 600-scoring peaks of NAC61 library using RSAT tool. (C) Functional enrichment analysis of NAC61 bound genes (assigned from –3kb to +100bp).

To define NAC61 high confident targets (HCTs), we then overlap the 1157 cv. ‘Thompson Seedless’ DEGs and the 2263 NAC61-bound unique genes (**Fig. 5A**). A total of 129 HCTs are thus identified (29 of which are in common with at least one of the GCNs; **Supplementary Dataset S5**). Interestingly, *NAC61* itself and other 15 annotated TFs are identified among the 78 HCTs, upregulated in *NAC61*-overexpressing plants, and further assigned to six clusters according to their expression profile in the global gene expression atlas (Fasoli *et al*., 2012; **Fig. 5B**). This allowed us to focus on the *NAC61* most closely correlated genes throughout the development of different grapevine organs.

**Figure 5:**
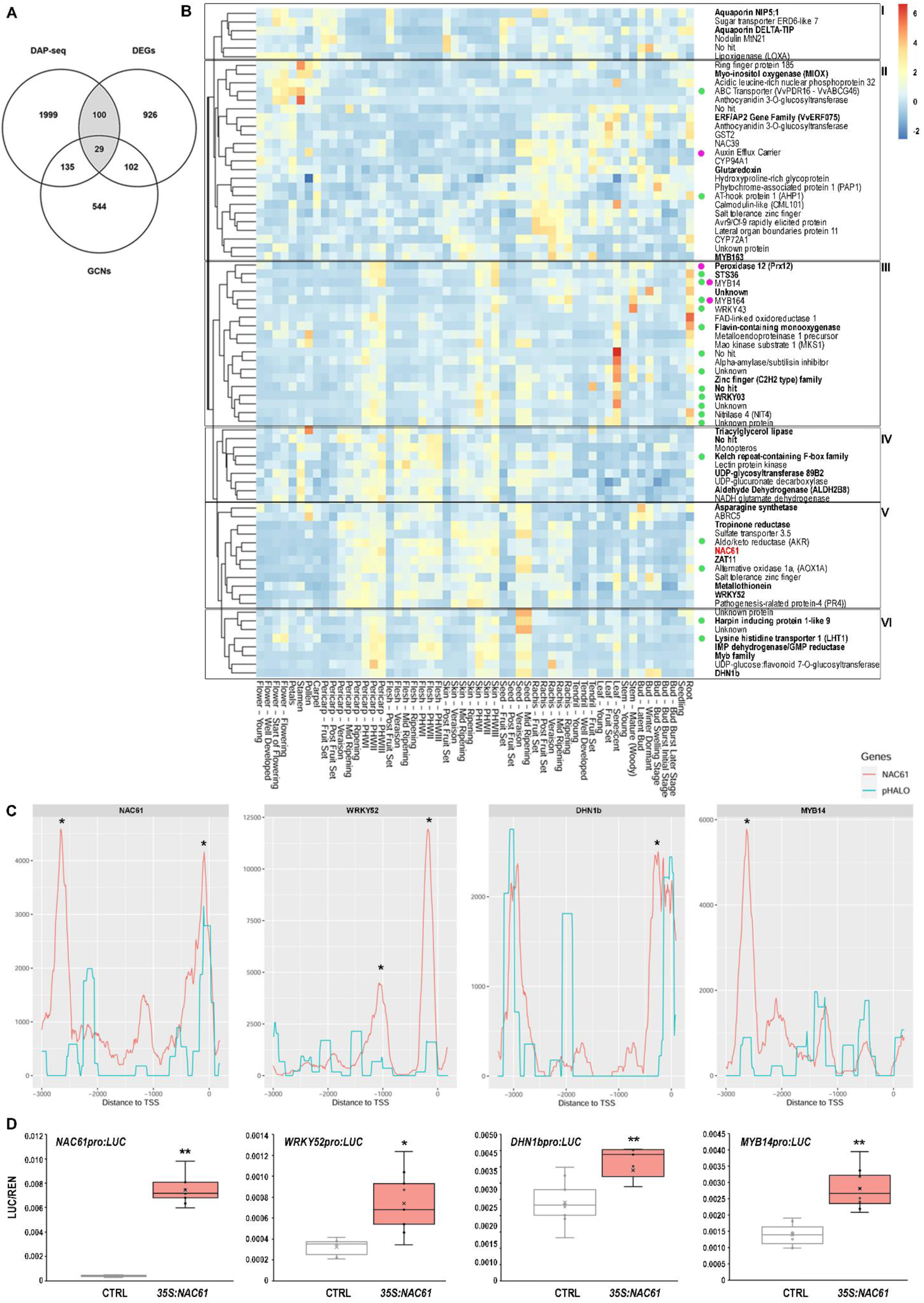
HCTs identification and validation. (A) Venn diagram showing the number of common genes between the DAP-seq bound genes (peaks in the -3kb and +100bp region), DEGs (FC ≥ 1.5 and adjust p-value <0.1) and GCNs (berry, leaf and TI datasets) (**Supplementary Dataset S5**). The NAC61 HCTs are in grey sections. (B) Heatmap representing the atlas expression (Fasoli *et al*., 2012) of the HCTs upregulated by the *NAC61* overexpression in the cv. ‘Thompson seedless’ leaves. The HCTs clusterization was performed through the Expression Atlases App (Corvina) within the VitViz platform (http://www.vitviz.tomsbiolab.com/), using the z-score data transformation and clustering by row. The 29 genes shared by the three datasets are highlighted in bold. Markers of the PHW process (Zenoni *et al*., 2016) and genes belonging to the STS GRN (Pilati *et al*., 2021) are marked with violet and green circle, respectively. (C) NAC61 DNA binding events shown as density plots and delimited between -3kb and +100bp from the TSS of *NAC61*, *WRKY52*, *DHN1b* and *MYB14*. The peaks were identified by GEM and were pointed out with their corresponding signal score in the proximal promoter regions. Asterisks (*) indicate the most significant peaks obtained by the DAP-seq analysis. (D) *NAC61*, *WRKY52*, *DHN1b* and *MYB14* promoter activation by NAC61 tested through the DLRA in infiltrated *Nicotiana benthamiana* leaves. *LUC* values are reported relative to the *REN* value. Each value represents the mean (± SD) of three biological replicates tested in technical replicate (*n=* 3). Asterisks indicate statistically significant differences (*, p < 0.05; **, p < 0.01; *t-*test).

Among the HCTs belonging to the *NAC61* cluster (Cluster V), we find several candidates putatively involved in abiotic and biotic stress responses such as two *zinc fingers*, an *aldo/keto reductase* (*AKR*), an *alternative oxidase* (*AOX1A*), the *pathogenesis-related protein 4* (*PR4*), encoding a chitinase, and *WRKY52*, recently reported as a *Botrytis cinerea* susceptibility gene (Wang *et al*., 2018). Albeit less closely correlated with NAC61 expression, other clusters include transcripts functionally associated to stress responses. In Cluster VI, we find a *harpin inducing protein 1-like 9*, a *lysine histidine transporter 1* (*LHT1*), and the previously described *DHN1b,* all highly expressed in berry during post-harvest dehydration and in seed after veraison. Moreover, the *aquaporins NIP5* and *DELTA-TIP,* the *nodulin MtN21* and a *lipooxigenase* (*LOXA*) are identified in Cluster I; a *myo-inositol oxygenase* (*MIOX*), a *glutathione S-transferase* (*GST2*), a *glutoredoxin,* a *calmodulin* (*CML101*), a *salt tolerance zing finger* and the *AVR9/CF-9 rapidly elicited protein a*re identified in Cluster II; an *aldehyde dehydrogenase* (*ALDH2B8*), the *NADH glutamate dehydrogenase,* the *monopteros, a triacylglycerol lipase* and a *kelch repeat-containing F-box family protein* are identified in Cluster IV. The above-described stilbenoid-related genes *STS36, MYB14, WRKY43,* and *WRKY03* are also found among the NAC61 HCTs. These genes, together with a *flavin-containing monooxygenase,* the *map kinase substrate 1* (*MKS1*), the *peroxidase 12* (*Prx12*), the *nitrilase 4* (*NIT4*) and the *MYB164*, are characterized by an increase of expression in berry skin during post-harvest dehydration, in senescing leaf and in root (Cluster III).

The NAC61-binding signal found in the *WRKY52, DHN1b, MYB14,* and *NAC61* promoters (**Fig. 5C)** is confirmed by DLRA, showing a significant activation by NAC61 (**Fig. 5D**).

### NAC61 participates in a NAC60-dependent regulatory network

*NAC61* was recently identified as a putative target of NAC60 (**Fig. 6A**; D’Inca *et al*., 2023) and shows a delayed expression pattern in comparison to that of *NAC60* in developing berries (**Fig. 1A**). Here, we demonstrate a significantly higher and lower *NAC61* expression level in grapevine leaves stably overexpressing *NAC60* and expressing the *NAC60* dominant repressor, respectively (**Fig. 6B**). Moreover, by performing DLRA, the ability of NAC60 to significantly transactivate the *NAC61* regulative region is demonstrated (**Fig. 6C**).

**Figure 6:**
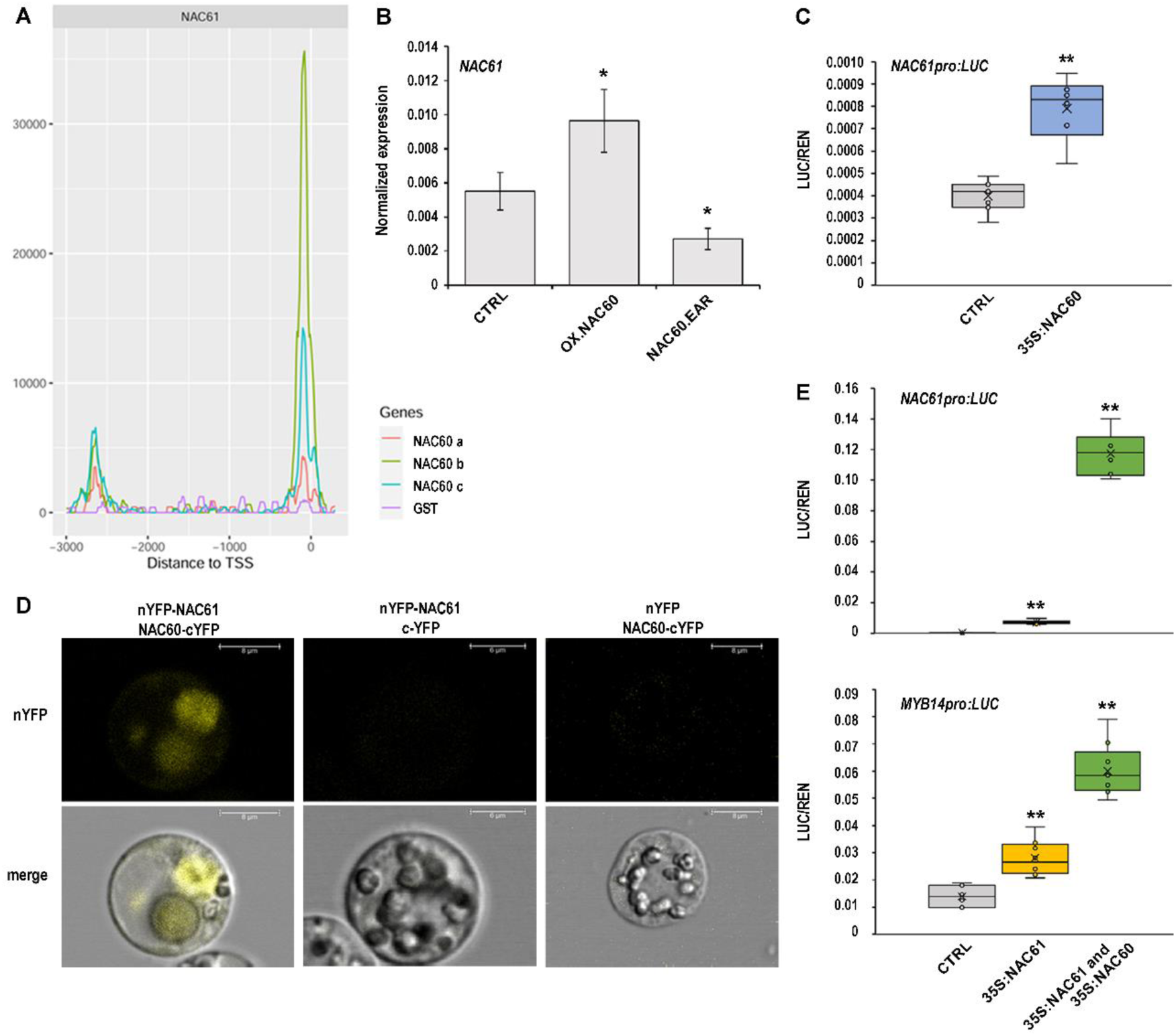
The NAC61-NAC60 regulatory complex regulates *NAC61* and *MYB14* activation. (A) NAC60 DNA binding events shown as density plots and delimited between -3kb and +100bp from the TSS of *NAC61.* The NAC60 binding motifs were searched in three different genomic libraries (a, berry gDNA; b and c are biological replicates of leaf gDNA; D’Incà *et al*., 2013) (B) *NAC61* expression level in grapevine leaves stably overexpressing *NAC60* (OX:NAC60) and expressing the *NAC60* dominant repressor (NAC60.EAR), determined by qPCR. Each value is relative to the *UBIQUITIN1* (*VIT_16s0098g01190*) and represents the mean (± SD) of three technical replicates. Asterisks indicate statistically significant differences (*, p < 0.05; *t-*test) in comparison to the control. (C) *NAC61* promoter transactivation by NAC60 tested by DLRA in infiltrated *Nicotiana benthamiana* leaves. *LUC* values are reported relative to the *REN* value. Each value represents the mean (± SD) of three biological replicates tested in technical replicate (*n=* 3). Asterisks indicate statistically significant differences (**, p < 0.01; *t-*test). (D) BiFC analysis in grapevine protoplasts showing NAC61/NAC60 protein interaction. Corresponding controls are also shown. Images show a representative case of YFP signal being detected in the cell nucleus, by using confocal laser scanning. (E) *NAC61* and *MYB14* promoter activation tested by DLRA in infiltrated *Nicotiana benthamiana* leaves. The single NAC61 (also reported in Fig. 5D) and combined NAC61-NAC60 activity were tested. *LUC* values are reported relative to the *REN* value. Each value represents the mean (± SD) of three biological replicates tested in technical replicate (*n=* 3). Asterisks indicate statistically significant differences (**, p < 0.01; *t-*test). The data reported in C and E have been performed in the same experiment and control values are therefore the same.

Both *NAC61* and *NAC60* were found as markers of the berry post-ripening phase (Zenoni *et al*., 2016) and *NAC60* is also found among the 92 common genes of the three *NAC61* GCNs datasets. Using BiFC assay, we demonstrate the physical interaction between the two NACs and reveal a heterocomplex localized into the nucleus (**Fig. 6D**). Accordingly, we show that NAC60 significantly increases NAC61 ability to activate the *MYB14* regulatory region (**Fig. 6E**). Indeed, we find that *MYB14* is induced 1.73 times by NAC61 alone, whereas the heterodimeric complex formed with NAC60 promotes a synergic action which reached a 4.29-time induction (**Fig. 6E**). The ability of NAC60 to significantly activate *MYB14* alone was previously demonstrated (D’Inca *et al*., 2023). We also demonstrate the NAC61 auto-activation (with a 5.59-time induction compared to the control; **Fig. 5C**), and the strong induction of its expression by the combined action of NAC61 and NAC60 (**Fig. 6E**).

By investigating the *NAC61* regulative region (-3.0 kb to the TSS), we find two binding sites for *A. thaliana* ANAC047 (**Supplementary Fig. S6**), coinciding with the already identified NAC60-binding site, and perfectly matching the NAC60-binding locations. We also find binding sites for RAP2.6, ABI3VP1/VRN1 and DEAR4, involved in ABA, osmotic, drought and salt stress responses (Wang *et al*., 2020a; Wang *et al*., 2020b; Zhang *et al*., 2020; Zhu *et al*., 2010b), for RRTF1 and RAP2.3, which mediate plant defense responses against *Botrytis cinerea* and other pathogens (León *et al*., 2020; Wang *et al*., 2020a), for ORA47, a regulator of general stress responses induced by methyl jasmonate (Zeng *et al*., 2022), and for DREB2C, AP2EREBP and G2like_tnt.At3g13040, involved in response to drought and dehydration (Dietz *et al*., 2010; Kizis *et al*., 2001; Lee *et al*., 2010; Riechmann and Meyerowitz, 1998; Wang *et al*., 2022), corroborating the *NAC61* role in abiotic and biotic stress responses (**Supplementary Fig. S6**).

### *NAC61* expression is enhanced by high temperature and *Botrytis cinerea* infection during berry post-harvest dehydration

Taking advantage of the experimental plan from a recent work (Shmuleviz *et al*., 2023), we analyze the expression of *NAC61* in grape berries subjected to post-harvest dehydration in two different temperature regimes. The results show that *NAC61* expression is significantly induced by high temperature, similarly to its HCT *MYB14* (**Fig. 7A**).

**Figure 7:**
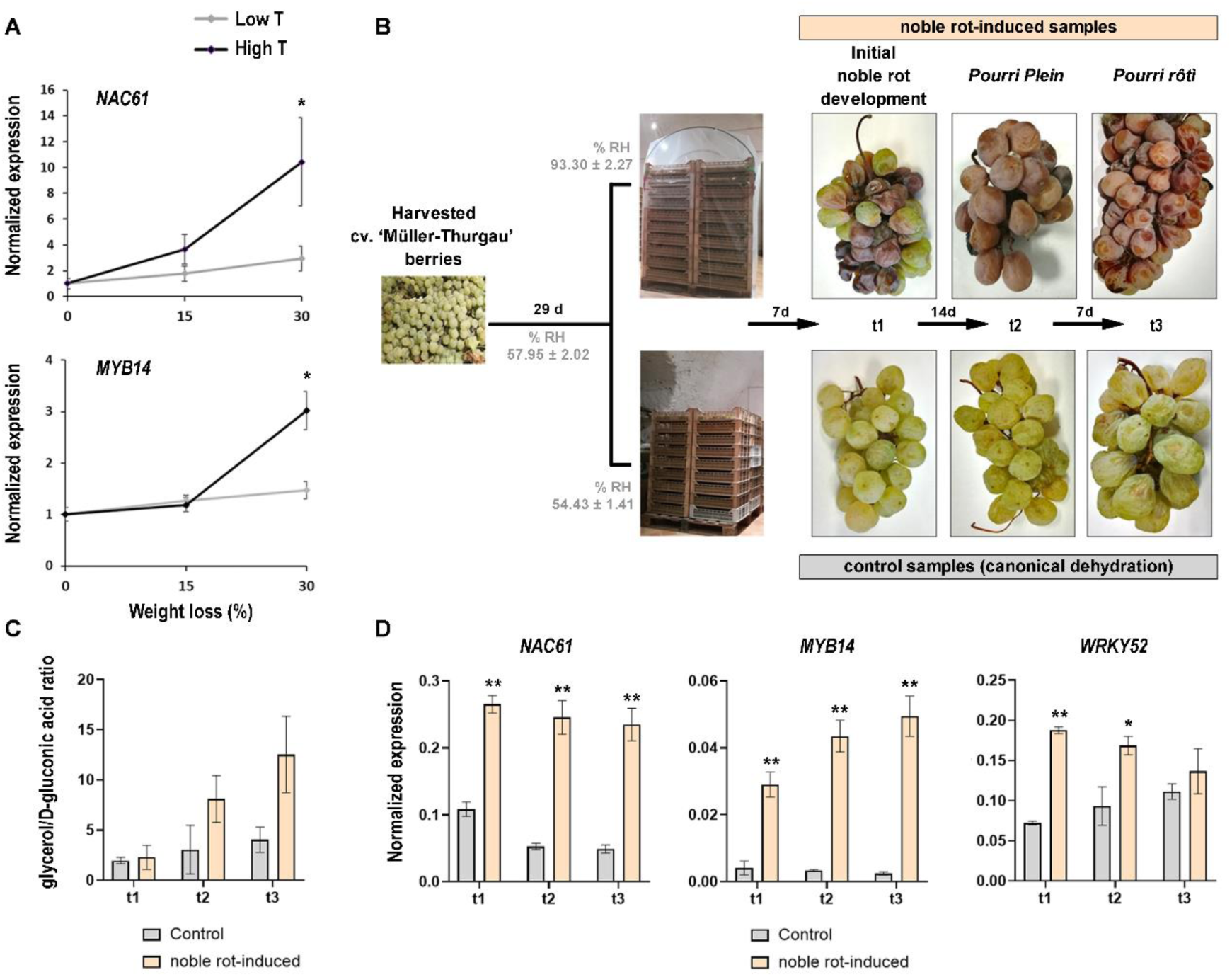
*NAC61* and target genes expression trend during post-harvest dehydration conducted in different conditions. (A) *NAC61* and *MYB14* expression level during post-harvest dehydration performed under high and low temperature conditions (Shmuleviz *et al*., 2023). Each value is relative to the *UBIQUITIN1* (*VIT_16s0098g01190*) and represents the mean (± SD) of three biological replicates. Asterisks indicate significant differences (*, p < 0.05; *t-*test). (B) Experimental plan for the noble rot induction. Berries of cv. ‘Müller Thurgau’ were collected at full maturity and put in a dehydrating room for 29 days to reach the 30% of weight loss. Then, half of the berries were covered to induce noble rot. The three stages (t0, t1 and t2) of infected and control berries collected for further analyses are represented. (C) Glycerol to D-gluconic acid ratio assessed as indicative of noble-rot development. Each value corresponds to the mean ± SD of three replicates. Asterisks indicate significant differences (*, p < 0.05; *t-*test). (D) *NAC61, MYB14* and *WRKY52* expression levels in noble rot-induced berries tested at different *B. cinerea* infection phases in cv. ‘Müller-Thurgau’ berries (noble-rot induced samples), compared to the control. Each value is relative to the *UBIQUITIN1* (*VIT_16s0098g01190*) and represents the mean (± SD) of three biological replicates. Asterisks indicate statistically significant differences (*, p < 0.05; **, p<0.01; t-test) of the noble rot-induced samples compared to the controls.

Because previous reports suggested that *NAC61* is induced during grape infection by *Botrytis cinerea*, in conditions of noble rot development but not of grey mold (Blanco-Ulate *et al*., 2015; Kelloniemi *et al*., 2015), we set up specific experimental conditions to induce noble rot in harvested berries (Negri *et al*., 2017), and analyze the expression of *NAC61* and some of its putative targets. Bunches of cv. ‘Müller-Thurgau’ were harvested at full maturity and placed in a ventilated dehydrating facility under controlled conditions (**Fig. 7B**). After 29 days of dehydration, when the soluble solid content reached ∼24°Brix, half of the bunches were covered with plastic film to naturally increase the relative humidity, required for noble rot induction (**Fig. 7B**). In these conditions, the dehydration and juice solute concentration are limited (**Supplementary Fig. S7**) and the first symptoms of noble rot appear on berries seven days after the coverage (t1). After 14 more days, infected berries reach the characteristic *pourri plein* stage (t2), and one week later, the clearly shriveled berries reach the *pourri rôtì* stage (t3) (**Fig. 7B;** Negri *et al*., 2017). The glycerol to d-gluconic acid ratio was assessed as indicative of noble-rot development (**Fig. 7C**) (Ribéreau-Gayon *et al*., 2006). The analysis of transcript levels by RT-qPCR, on control and noble rot-induced berries, shows a strong responsiveness of *NAC61*, and the NAC61 targets *MYB14* and *WRKY52* during noble rot development (**Fig. 7D**).

## DISCUSSION

### NAC61 controls the stilbenoid biosynthetic pathway as a conserved feature of late-and post-ripening

Stilbenoids are a group of polyphenols synthetized by STSs in response to biotic, abiotic and developmental cues. *STSs* are also expressed in the absence of external stimuli in a tissue and cultivar-dependent manner (Eisenmann *et al*., 2019; Gatto *et al*., 2008; Jean-Denis *et al*., 2006; Pezet *et al*., 2003; Versari *et al*., 2001). In grapevine, *STSs* represent a large gene family, encompassing 41 isoforms (Vannozzi *et al*., 2012), most of which are regulated by Subgroup 2 R2R3-MYB transcription factors (i.e., MYB13/MYB14/MYB15; Orduña *et al*., 2022). Among the 12 transiently-activated *STSs* in *NAC61*-overexpressing leaves (**Fig. 3B**), *STS36* (Huang *et al*., 2018) is identified as a candidate NAC61 HCT. Additionally, our results demonstrate that NAC61 binds to, and directly regulates, *MYB14*, together with *WRKY03* and *WRKY43*, all of which cooperate in enhancing *STSs* expression (Vannozzi *et al*., 2018). These results allow to place NAC61 on a high-hierarchical position for the regulation of stilbene synthesis. Also corroborating this NAC61-dependent transcriptional cascade, 117 out of 530 genes upregulated by *NAC61-*overexpression belong to the recently described *STSs* GRN displaying more than one thousand structural and regulatory genes potentially involved in stilbenoid metabolism (**Fig. 3B, Supplementary Dataset S2)** (Pilati *et al*., 2021). Together with the above-described HCTs (*STS36*, *MYB14*, *WRKY03* and *WRKY43*), we identify two other R2R3-MYB members, *MYB163* and *MYB164*, previously associated with the phenylpropanoid pathway (Wong *et al*., 2016).

In grapevine, LACs have been proposed to control the oxidative polymerization of both monomeric stilbenes and monolignols, producing viniferins and lignin, respectively (Keylor *et al*., 2015). *Vitis vinifera* LAC family members were recently assigned to the stilbenoid-and lignin-related subgroups based on sequence similarity and co-expression (Pilati *et al*., 2021). Interestingly, among the 33 *LACs* being upregulated in *NAC61*-overexpressing grapevine plants, 23 belong to the stilbenoid subgroup, while none of these belong in lignin-related subgroup. Nevertheless, the absence of *LAC* genes among the HCTs indicates that NAC61-mediated *LAC* regulation may occur indirectly, likely through MYB14 as shown by Orduna *et al*. (2022).

Among NAC61 HCTs, we also find the *AKR,* the *AOX1A*, the *AT-hook protein 1 (AHP1)*, a *flavin-containing monooxygenase*, the *harpin inducing protein 1-like 9*, the *kelch repeat-containing f-box family protein*, and the *LHT1*, to be present in the *STSs* GRN (**Supplementary Dataset S5)**. Despite the role of these genes in stilbenoid metabolism, it is not clear and further investigations are needed, their belonging to the NAC61 HCTs strongly support the master regulatory role of NAC61 in stilbenes synthesis and modification. The accumulation of stilbenoids hallmark late-and post-ripening stages, thus the genes related to their synthesis, can be considered true markers of these developmental transitions. Consistently, 34 out of the 75 total PHW molecular markers defined by Zenoni *et al*. (2016) are upregulated by *NAC61*-overexpression, including *JAZ4,* eight STSs, six *LACs,* the *dirigent protein DIR16*, four *nitrilases*, an *osmotin*, the *Prx12* and a *pathogenesis-related protein*, in addition to the previously mentioned *MYB14, MYB164, WRKY03,* and *WRKY43* (**Fig. 3B**; **Supplementary Dataset S2)**.

### NAC61 is responsive to osmotic stress

During late-and post-ripening stages, grape berries are subjected to a progressive increase of solute concentration, determining a severe osmotic stress. Several evidences, arising from our study, strongly indicate NAC61 involvement in osmotic stress responses: (i) the close correlation between sugar concentration and *NAC61* expression, (ii) the earlier activation of *NAC61* in berry flesh, where sugars and other metabolites accumulate (in comparison to the skin), (iii) the higher RWC in *N. benthamiana NAC61*-expressing leaves in comparison to the control, and (iv) the identification of NAC61 HCTs potentially involved in osmotic stress response through different strategies/mechanisms.

Four zinc finger protein-coding genes are found among HCTs: two *salt tolerance zinc fingers*, a *zinc finger C2H2 type* and the *C2H2-type zinc finger ZAT11*, belonging to the branch of the C2H2 family containing the ZAT domain, whose role in response to abiotic stresses in several plant species has been widely described (Liu *et al*., 2022). Interestingly, *cis*-regulatory motifs of ZAT TFs were found in *dehydrin* (*DHN*) genes recently characterized in *Brachypodium* grasses (Decena *et al*., 2021). *DHNs* are ubiquitously expressed in low intracellular water content period (Liu *et al*., 2017; Smith and Graether, 2022; Yang *et al*., 2012) and their role in coping with osmotic stress has been demonstrated in several species (Tiwari and Chakrabarty, 2021). In our study, we identify and validate *DHN1b* as a NAC61 target. *DHN1b* is upregulated in withering grape berries (Zamboni *et al*., 2008) and in leaves and berries subjected to water stress (Savoi *et al*., 2017; Xiao and Nassuth, 2006). Further investigation to elucidate a putative NAC61 and ZAT11 cooperation in regulating *DHN1b* expression in osmotic stress conditions would deserve attention in future studies.

NAC61 HCTs include also *LHT1,* a major candidate for root acquisition of aspartate and a transporter of aspartate, asparagine and glutamate in rice (Guo *et al*., 2020). In grapevine, the induction of a *lysine histidine transporter* and other amino acid transporters in salt-stressed plants has been observed (Aydemir *et al*., 2020). This finding, together with the increased RWC in the *NAC61-*overexpressing *N. benthamiana* leaves, suggests that NAC61 could exert its role in water/osmotic stress response by regulating amino acid accumulation. Accordingly, an *asparagine synthase* is the most strongly upregulated HCT by *NAC61* overexpression. This enzyme catalyzes the synthesis of asparagine from aspartate and asparagine is one of the most represented amino acids in grape berries (Bouloumpasi *et al*., 2015). Together these two amino acids are widely described as a drought-responsive metabolites (Han *et al*., 2021). In addition, a *glutamate dehydrogenase* (*GDH*), which catalyzes the oxidative deamination of glutamate to generate α-ketoglutarate (Sullivan *et al*., 2015) thus providing the carbon for *de novo* synthesis of aspartate, is also identified among NAC61 HCTs.

Osmotic stress responses could also involve auxin metabolism that in turn may contribute to drought tolerance through stomata closure regulation. Interestingly, an *auxin efflux carrier* and the *ARF TF Monopteros*, which inhibits stomatal development (Zhang *et al*., 2014), is found among NAC61 HCTs. Similarly, the downregulation of the HCTs *histidine phosphotransfer AHP4*, whose knock-out narrows stomatal apertures, heightens leaf temperatures during water stress and increases leaf RWC (Ha *et al*., 2022) and the *phosphatase PP2CA/AHG3*, whose repression activates ABA-mediated signaling pathway leading to stomatal closure and water retention (Jung *et al*., 2020), may contribute to NAC61 function in osmotic stress responses. Interestingly, hypoxia-related genes are found among *NAC61* induced DEGs, such as a *Hypoxia-responsive gene* and three dehydration-responsive proteins (*RD22*), indicating a decrease in oxygen supply likely due to water saturation of the apoplast (van den Dries *et al*., 2013).

The remodeling of lipid composition, to maintain the fluidity and stability of cell membranes, is another change adopted by plants to respond to osmotic stress. In this regard, among HCTs, we find a *triacylglycerol lipase,* whose induction in grape berry under water deficit was already reported (Savoi *et al*., 2017). During prolonged drought stress, the membranes could be subjected to degradative processes, not only due to lipolytic activity but also peroxidative activity. In this context, we could hypothesize an involvement of two NAC61 HCTs, *Prx12* and *LOXA,* both also known to be involved in pathogen responses. The activity of these two enzymes could be related to the cell death observed in *NAC61*-overexpressing leaves three days post-agroinfiltration.

### NAC61 regulates redox state and defense genes during noble rot development

The significant increase of H_2_O_2_ observed in *NAC61*-overexpressing *N. benthamiana* leaves and the NAC61-mediated upregulation of five *Peroxidases (Prxs),* including the HCT encoding *Prx12,* may account for apoplastic ROS production (Survila *et al*., 2016). This strongly suggests a direct involvement of NAC61 in ROS accumulation. Interestingly, *Prxs* are a well-known class of PR proteins induced in host plant tissues by pathogen infection (Lüthje and Martinez-Cortes, 2018), suggesting also a direct involvement of NAC61 in biotic stress responses. Accordingly, besides genes related to stilbenoid synthesis and osmotic stress that may account for grapevine defense against pathogens (Yang *et al*., 2012), we also find several other HCTs related to biotic stress responses, such as *PR4*, the biotic stress-responsive calmodulin-like *CML101* (Vandelle *et al*., 2018), the *harpin inducing protein 1-like 9* and *AKR* also identified as NAC60 targets (D’Inca *et al*., 2023), an *Avr9/Cf-9 rapidly elicited protein*, *LOXA* and *MKS1*.

The upregulation of genes encoding pathogen-related proteins has been previously evidenced in healthy berries during PHW as part of a general response to biotic stresses (Zenoni *et al*., 2016). We could then hypothesize that NAC61 is involved in ROS metabolism/homeostasis, on the one hand, by regulating the expression of genes involved in defense, while also activating mechanisms for ROS detoxification. Indeed, several genes involved in ROS scavenging/detoxification are found as HCTs, such *MIOX,* a *flavin-containing monooxygenase, AOX1A,* an *aldehyde dehydrogenase (ALDH288*), a *glutaredoxin*, a *GDH* and *GST2* (Munir *et al*., 2020; Vanlerberghe *et al*., 2020; Wang *et al*., 2023; Zhang *et al*., 2012), making very intricated the involvement of NAC61 in ROS metabolism.

ROS are also crucial signals for the induction of the hypersensitive response, a programmed cell death process that facilitates plant infection by necrotrophic pathogens (Soosaar *et al*., 2005), including *Botrytis cinerea* (Govrin and Levine, 2000), responsible for grey mold in grapes. However, in particular conditions, the fungus leads to the development of noble rot, which promotes favorable changes in grape berries thanks to a weaker (or even controlled) infection. The strong induction of *NAC61* in botrytized berries, together with the HCT *WRKY52,* recently characterized as a grapevine susceptibility gene of *B. cinerea* (Wang *et al*., 2018), indicates a possible direct involvement of *NAC61* in providing favorable conditions for noble rot development. This hypothesis is further supported by the identification among HCTs of *MKS1,* whose overexpression in *A. thaliana* was shown to increase susceptibility to *B. cinerea* (Petersen *et al*., 2010) and to promote the upregulation of *Prx 12* specifically during noble rot (Blanco-Ulate *et al*., 2015; Lovato *et al*., 2019). Moreover, the *VqSTS36* was shown to enhance *B. cinerea* susceptibility in *A. thaliana* and Tomato (Huang et al., 2018). On the other hand, NAC61 could also contribute to a weaker infection by simultaneously mitigating the favorable conditions for *B. cinerea* growth through the regulation of *PR4*, encoding a chitinase, which could inhibit the growth of fungal hyphae (Grover, 2012), and *LOXA,* involved in the biosynthesis of jasmonate, known to mediate defense against necrotrophic pathogens (Antico *et al*., 2012). Furthermore, in *A. thaliana* the overexpression of JAZ8 represses defense responses against *B. cinerea* through its interaction with AtWRKY75 (Chen *et al*., 2020). Considering that *WRKY52* is one of the closest homologues of *AtWRKY75* (Vannozzi *et al*., 2018), and that two *JAZs*, namely *JAZ2* and *JAZ4*, are upregulated by NAC61, a similar mechanism that allows noble rot development could be suggested in grape berries.

Interestingly, *Botrytis elliptica* infection of *Lilium regale* downregulates *miR164* (Gao *et al*., 2017) and the transient *miR164f* overexpression in apple leaves enhances their susceptibility to *Alternaria alternata* AP, possibly due to the downregulation of a *NAC* TF (Zhou *et al*., 2023). Moreover, several NAC members, belonging to the same clade of NAC61, are post-transcriptionally regulated by the *miR164* family (Kim *et al*., 2009; Lira *et al*., 2017; Sun *et al*., 2012). We could therefore assume a scenario in which *B. cinerea* may affect *miR164* expression in late berry ripening stage to allow *NAC61* transcript level increase. Similarly, low temperature storage conditions, which ultimately lead to fruit senescence delay, repress the strawberry genes *FaNAC087* and *FaNAC038* due to an increase of their negative regulator *miR164* (Li *et al*., 2017; Xu *et al*., 2013). Since *NAC61* also shows a lower expression in berries experiencing post-harvest dehydration under low temperature conditions, the control by *miR164* might be conserved in grape.

### NAC-dependent transcriptional network behind berry aging and stress responses

The GCNs obtained from different grapevine organs reveal a strong correlation between *NAC61* and many genes previously associated with late-and post-ripening developmental stages (**Supplementary Dataset S1**). Consistently, NAC61 HCTs include several genes involved in stilbenoid metabolism, as well as osmotic and biotic stress responses, which characterize these late processes (**Fig. 8**), thus suggesting NAC61 as a key regulator triggering the molecular mechanisms controlling ripening progression.

**Figure 8:**
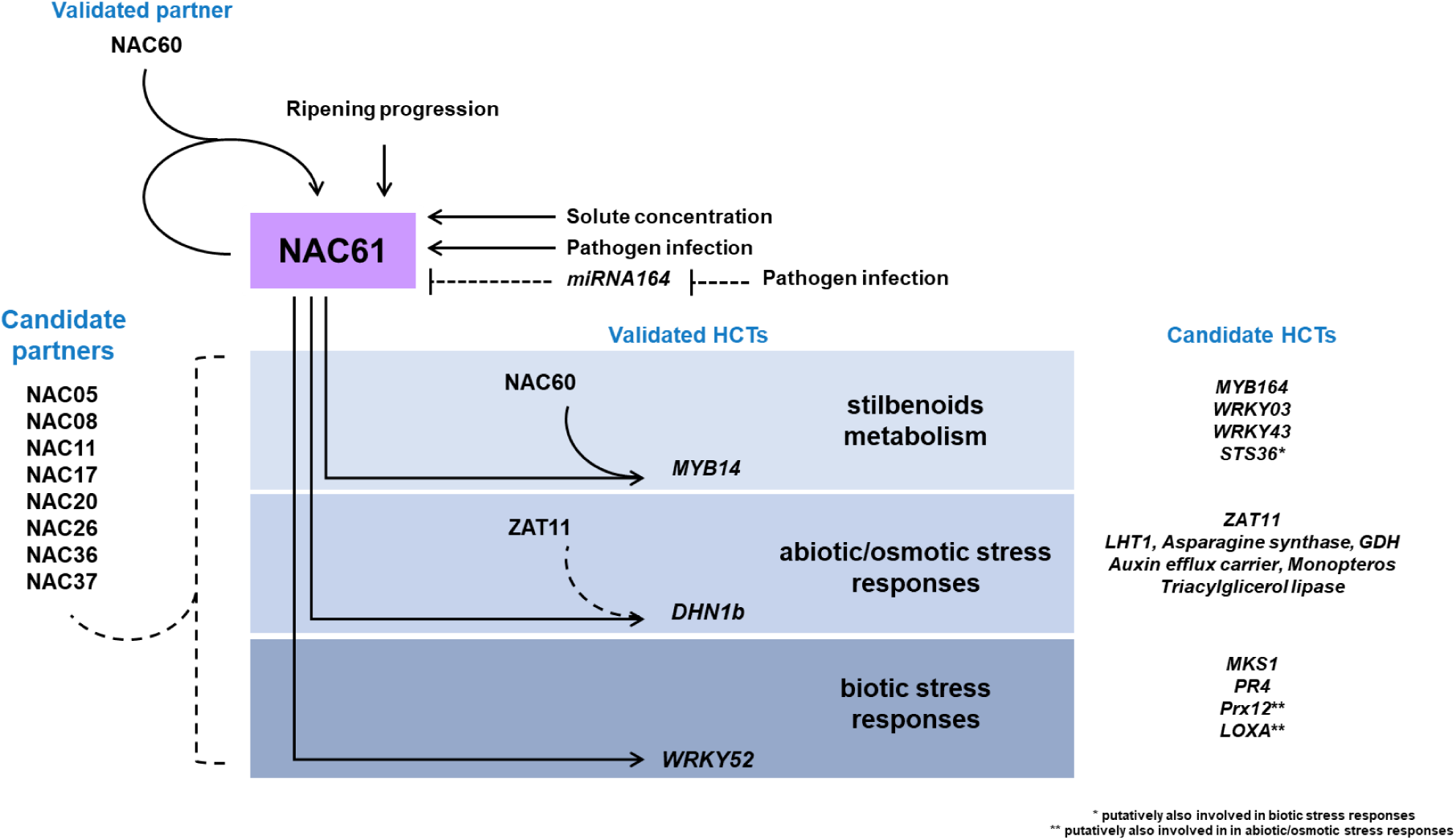
Proposed model of NAC61 involving regulatory network.

Interestingly, NAC60, which belongs to *NAC61* GCN, not only activates the expression of NAC61 but also forms heterodimers with it, providing a mechanism for the transactivation of their common target *MYB14*. Moreover, NAC61 self-activates, though a NAC60-NAC61 heterodimer that greatly increases *NAC61* expression, indicating that NAC61 participates in the NAC60-dependent regulatory network (**Fig. 8**).

Besides, other *NACs* are co-expressed with *NAC61,* thus representing putative partners. These include *NAC11*, previously described as a berry ‘switch gene’ by Massonnet *et al*. (2017), *NAC17*, involved in salinity and drought stress responses (Ju *et al*., 2020) and *NAC26*, proposed as a determinant of berry size variation (Muñoz-Espinoza *et al*., 2020; Tello *et al*., 2015) and a regulator of seed and fruit development through the interaction with MADS9 (Zhang *et al*., 2021).

We therefore propose the existence of a NAC60-dependent NAC network governing grape berry development. Further lines of research should focus on the characterization of NAC61 downstream target genes and interacting proteins to elucidate the molecular mechanisms underlying berry aging and stress responses.

## SUPPLEMENTARY DATA

**Supplementary Figure S1.** *NAC61* expression pattern during berry development. **Supplementary Figure S2.** Gene Ontology (GO) enrichment of the *NAC61* co-expressed genes. **Supplementary Figure S3.** NAC61-containing cluster in the NACs phylogenetic tree.

**Supplementary Figure S4.** *NAC61* overexpression in cv. ‘Thompson Seedless’ grapevine leaves.

**Supplementary Figure S5.** NAC61 binding motif discovery analysis and motif comparison with published *A. thaliana* datasets.

**Supplementary Figure S6.** *NAC61* regulative region analysis for *A. thaliana* ANAC047 and stress-related proteins *cis*-elements performed with the RSAT software.

**Supplementary Figure S7.** Juice solute concentration in noble rot-induced and control grape berries of cv “Muller Thurgau”.

**Supplementary Table S1.** List of used primers.

**Supplementary Dataset S1.** *NAC61* co-expressed genes. The GCNs were obtained separately by

**Supplementary Dataset S2.** Transcriptomic analysis of *NAC61-*overexpressing and control cv. ‘Thompson Seedless’ leaves.

**Supplementary Dataset S3.** Gene category MapMan distribution and enrichment analysis of DEGs.

**Supplementary Dataset S4.** NAC61 DAP-seq bound genes.

**Supplementary Dataset S5.** List of defined HCTs and detail of genes grouped in Fig. 5A.

### Abbreviations

(DAB): 3,3’-Diaminobenzidine
(ABA): Abscisic Acid
(BiFC): Bimolecular Fluorescence Complementation Assay
(CDS): Coding Sequence
(DEGs): Control (CTRL), Differentially Expressed Genes
(DAP-seq): DNA Affinity Purification and sequencing
(DLRA): Dual Luciferase Reporter Assay
(LUC): *Firefly* Luciferase
(FC): Fold Change
(GCNs): Gene Co-expression Networks
(GRN): Gene Regulatory Network
(HCTs): High Confident Targets
(PHW): Post Harvest Withering
(ROS): Reactive Oxygen species
(RT-qPCR): Real Time quantitative Polymerase Chain Reaction
(RWC): Relative Water Content
(REN): *Renilla reniformis* Luciferase
(TF): Tissue-independent (TI), Transcription Factor
(TSS): Transcription Start Site
(YFP): Yellow Fluorescent Protein.

## Acknowledgments

The authors thank Mario Pezzotti (University of Verona, Italy) for supporting this study and for critical discussions and Ron Shmuleviz for providing the cv. ‘Corvina’ mature and low/high temperature dried berries cDNA. The authors would like to thank the Genomics and Transcriptomics platform of Centro Piattaforme Tecnologiche (CPT), University of Verona. All the bioinformatic analyses were performed on the HPC cluster Garnatxa at Institute for Integrative Systems Biology (I2SysBio).

## Author contributions

C.F., G.B.T., A.A. and S.Z. designed the research; C.F., D.D., O.B. and A.A performed the research; C.F., L.O., J.T.M., A.A. and S.Z. analyzed data; C.F., L.O. and J.T.M. contributed with analytic and computational tools and analyzed data; C.F., J.T.M., E.V., G.B.T., A.A. and S.Z. wrote the paper. All authors have read and agreed to the published version of the manuscript.

## Conflict of interest

The authors have no conflict of interest to declare.

## Funding

This work was supported by Grant Ricerca di Base ‘Definition of master regulator genes of fruit ripening in grapevine’, University of Verona, awarded to S.Z., by PRIN 2017 ‘Regulation of gene expression in grapevine: analysis of genetic and epigenetic determinants’ awarded to Mario Pezzotti, by grants PID2021-128865NB-I00 and RYC-2017-23645 awarded to J.T.M. and PRE2019-088044 fellowship awarded to L.O. from the Ministerio de Ciencia, Innovación y Universidades (MCIU, Spain), Agencia Estatal de Investigación (AEI, Spain), and Fondo Europeo de Desarrollo Regional (FEDER, European Union). This article is based upon work from COST Action CA 17111 INTEGRAPE, supported by COST (European Cooperation in Science and Technology).

## Data availability

Microarray data for the transient expression experiments on *V. vinifera* cv. ‘Thompson Seedless’ are available at GEO under the accession ‘GSE232165’. DAP-seq raw data has been submitted to GEO, including metadata of samples and conducted analysis (bioinformatic parameters) according to the FAIR principles, under the accession ‘GSE230185’. DAP-seq results on NAC61 can be visualized in the DAPBrowse tool available at the Vitis Visualization Platform (http;//www.vitviz.tomsbiolab.com/). The role for *NAC61* has been deposited in the Gene Reference Catalogue found at the Grape Genomics Encyclopedia portal (http://grapedia.org/).

## Notes

### Competing Interest Statement

The authors have declared no competing interest.

