## Supplementary Figures and Table for "NAC61 regulates late-and post-ripening associated processes in grapes involving a NAC60-dependent regulatory network"

**Supplementary Figure S1.** *NAC61* expression pattern during berry development. (A) *NAC61* expression profile in 10 different grapevine varieties at two pre- (pea and touch) and two post- (soft and harvest) veraison developmental stages (Massonnet *et al.*, 2017). (B) *NAC61* expression profile in cv. 'Cabernet Sauvignon' and 'Pinot noir' berries sampled every 10 days from fruit set to ripening (Fasoli *et al.*, 2018). (C) *NAC61* cluster of gene expression in the Genotype x Environment (GxE) dataset and box plot of Variable Importance Measure (VIM) used to characterize the relationship between the cluster and the experimental conditions (Dal Santo *et al.*, 2018). (D) *NAC61* expression trend in cv. 'Corvina' berries during traditional long and forced short post-harvest dehydration processes (Zenoni *et al.*, 2020). Weight loss percentage is specified for each point.

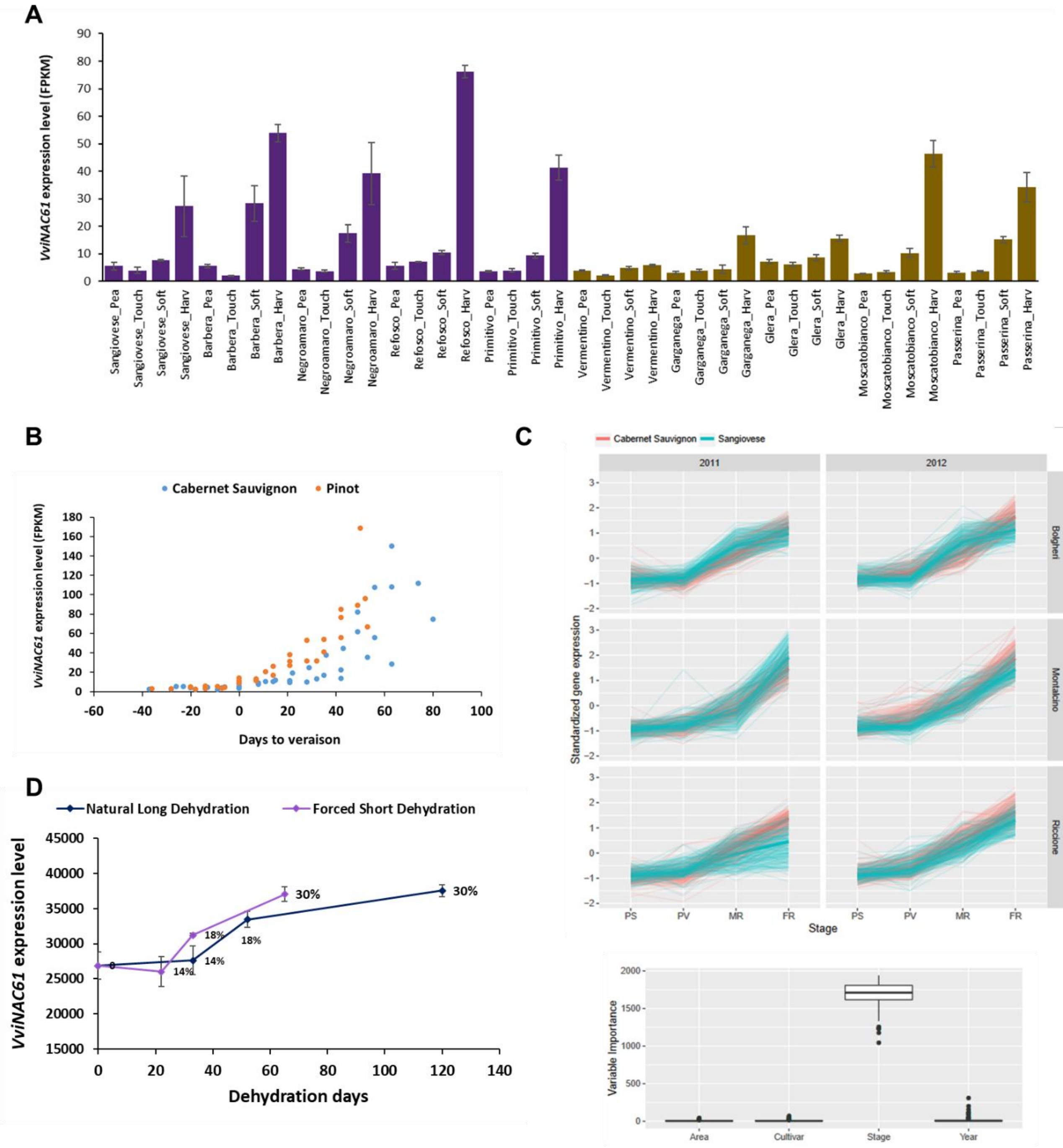
A
VviNAC61 expression level (FPKM)
90
80
70
60
50
40
30
20
10
0
Sangiovese_Pea
Sangiovese_Touch
Sangiovese_Soft
Sangiovese_Harv
Barbera_Pea
Barbera_Touch
Barbera_Soft
Barbera_Harv
Negroamaro_Pea
Negroamaro_Touch
Negroamaro_Soft
Negroamaro_Harv
Refosco_Pea
Refosco_Touch
Refosco_Soft
Refosco_Harv
Primitivo_Pea
Primitivo_Touch
Primitivo_Soft
Primitivo_Harv
Vermentino_Pea
Vermentino_Touch
Vermentino_Soft
Vermentino_Harv
Garganega_Pea
Garganega_Touch
Garganega_Soft
Garganega_Harv
Glera_Pea
Glera_Touch
Glera_Soft
Glera_Harv
Moscatobianco_Pea
Moscatobianco_Touch
Moscatobianco_Soft
Moscatobianco_Harv
Passerina_Pea
Passerina_Touch
Passerina_Soft
Passerina_Harv
B
Cabernet Sauvignon
Pinot
VviNAC61 expression level (FPKM)
180
160
140
120
100
80
60
40
20
0
-60
-40
-20
0
20
40
60
80
100
Days to veraison
C
Cabernet Sauvignon
Sangiovese
2011
2012
Bolgheri
Montalcino
Riccione
Standardized gene expression
3
2
1
0
-1
-2
PS
PV
MR
FR
Stage
2000
1500
1000
500
0
Variable Importance
Area
Cultivar
Stage
Year
D
Natural Long Dehydration
Forced Short Dehydration
VviNAC61 expression level
45000
40000
35000
30000
25000
20000
15000
0
20
40
60
80
100
120
140
Dehydration days
0
14%
14%
18%
18%
30%
30%

**Supplementary Figure S2.** Gene Ontology (GO) enrichment of the *NAC61* co-expressed genes. The GO enrichment analysis of the 810 genes co-expressed with *NAC61* (Supplementary Dataset S1) was performed by using the ShinyGO v.0.741 software (Ge *et al.*, 2020) with a False Discovery Rate (FDR) cutoff of 0.001.

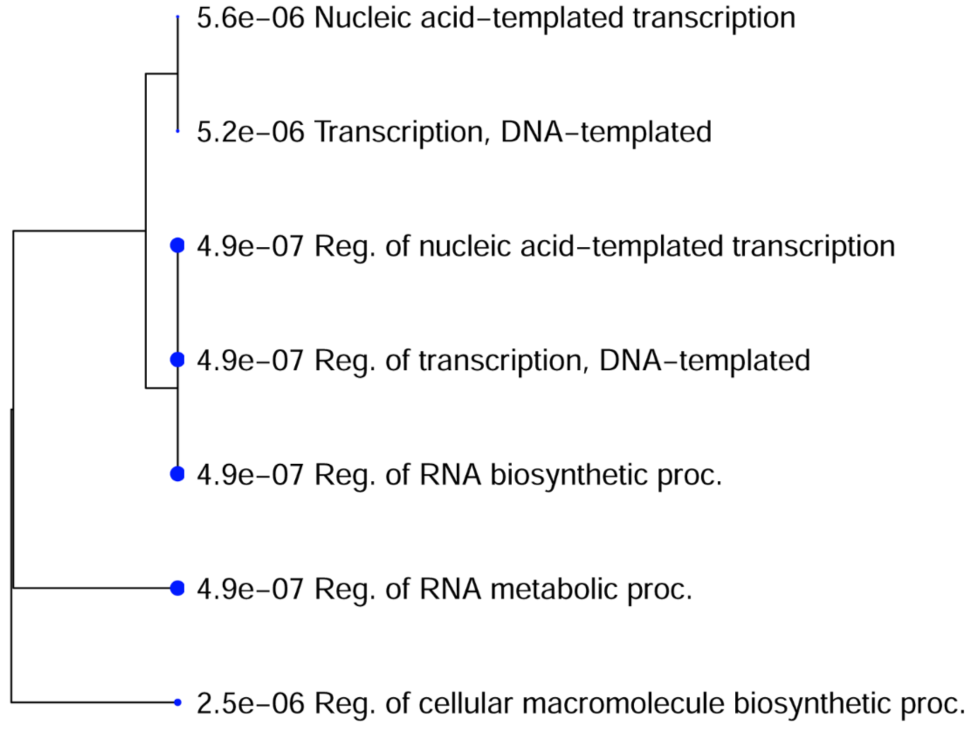

**Supplementary Figure S3.** NAC61-containing cluster in the NACs phylogenetic tree. Phylogenetic relationships of different plant species NAC genes were previously investigated (https://tomsbiolab.com/wp-content/uploads/2021/10/Fig.-S4.png; D’Incà *et al.*, 2023) and the NAC61-containing cluster is here reported. Red rows indicate ANAC046, the VvNAC61 closest homolog, VviNAC61 and VviNAC33.

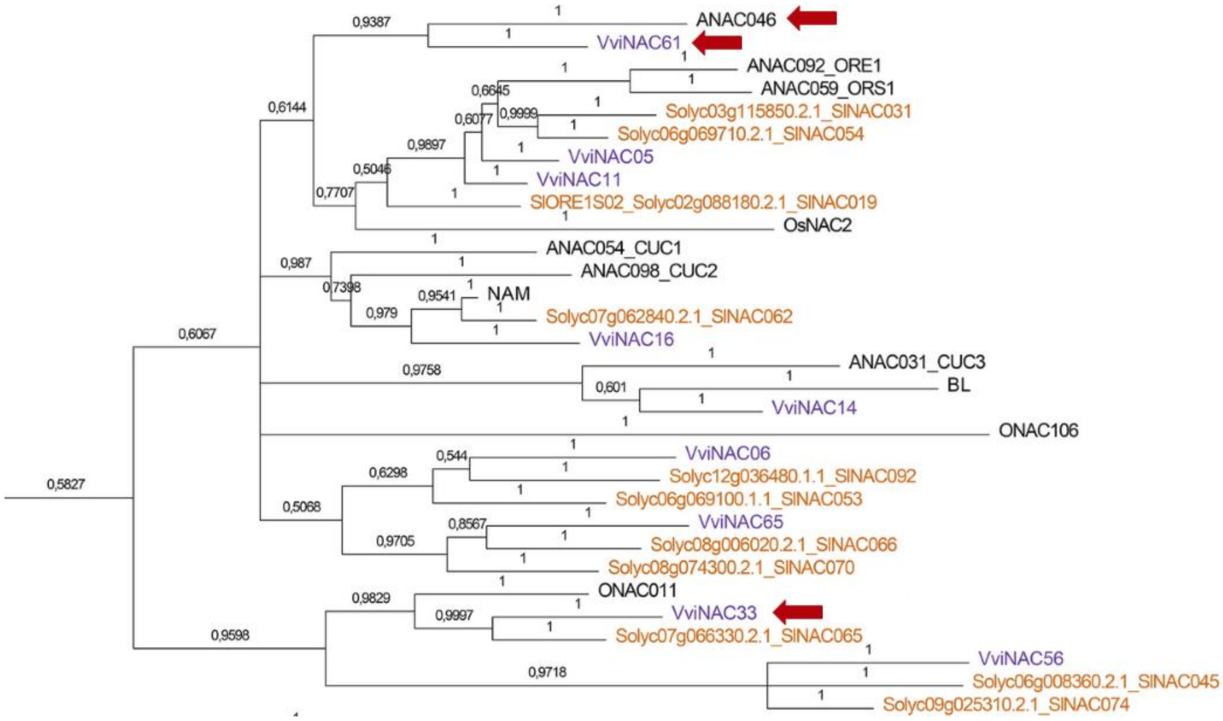

**Supplementary Figure S4.** *NAC61* overexpression in cv. 'Thompson Seedless' grapevine leaves. (A) The significant *NAC61* transient overexpression (*, p<0.01; t-test) in comparison to the control was validated by RT-qPCR. Each value corresponds to the mean ± SD of three technical replicates relative to the *UBIQUITIN1*. (B) Expression level of *MYB14* and of a *laccase* gene (*VIT_18s0001g01280*) in *NAC61* overexpressing and in control lines determined by RT-qPCR. Each value corresponds to the mean ± SD of three biological and three technical replicates relative to the *UBIQUITIN1*. Asterisks indicate statistically significant differences (*, p< 0.01; t-test).

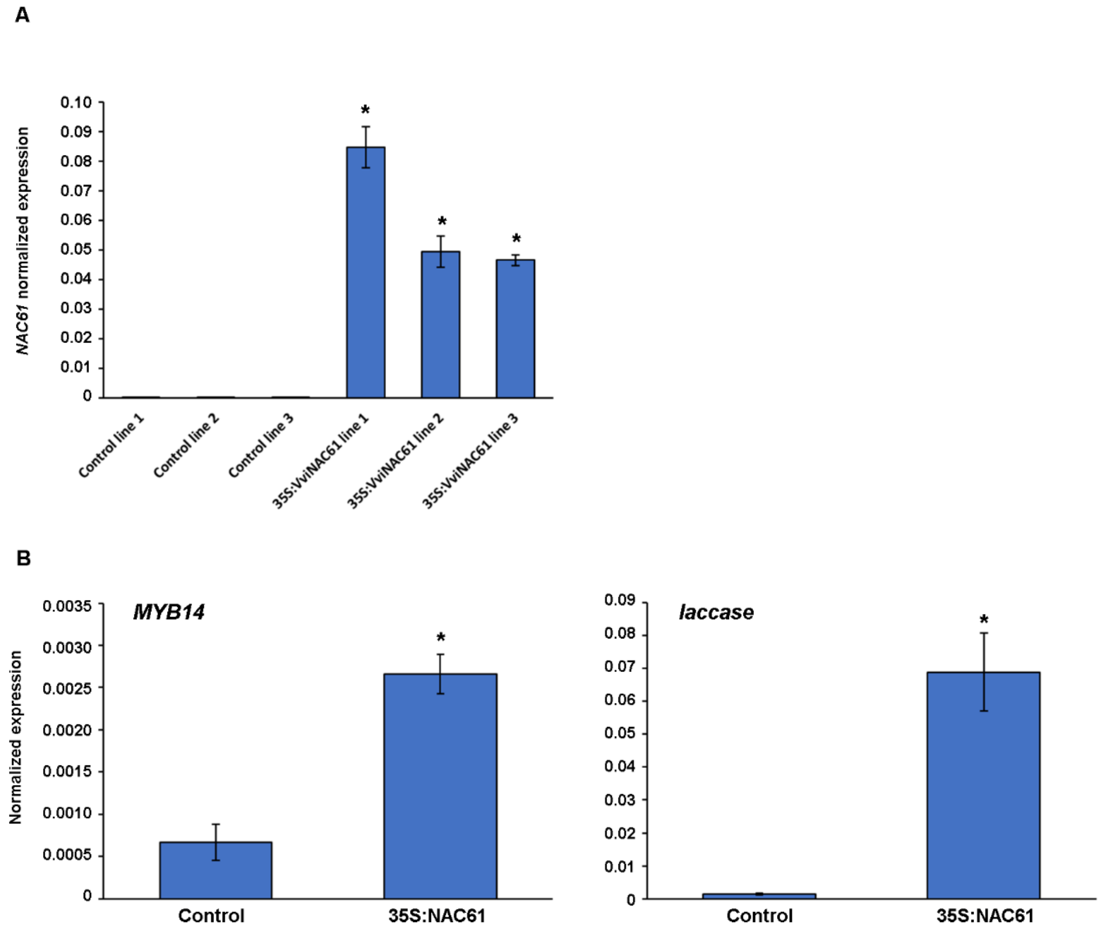

**Supplementary Figure S5.** NAC61 binding motif discovery analysis and motif comparison with published *A. thaliana* datasets.

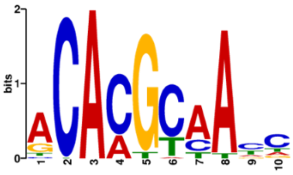


k-mer sig = **55.08**; evalue = **8.3e-56**

**ArabidopsisPBM**

| Name | ID | Strand | Nb overlap columns | % aligned | Pearson correlation | Normalized cor |
|---|---|---|---|---|---|---|
| 7195_ANAC58_ArabidopsisPBM_20140210_ | 7195_ANAC58_ArabidopsisPBM_20140210_ | D | 10 | 0.6250 | 0.943 | 0.590 |
| 7192_ANAC46_ArabidopsisPBM_20140210_ | 7192_ANAC46_ArabidopsisPBM_20140210_ | D | 10 | 0.6250 | 0.901 | 0.563 |
| 7193_ANAC55_ArabidopsisPBM_20140210_ | 7193_ANAC55_ArabidopsisPBM_20140210_ | D | 10 | 0.6250 | 0.860 | 0.537 |

Total matches = 3

**Athamap**

| Name | ID | Strand | Nb overlap columns | % aligned | Pearson correlation | Normalized cor |
|---|---|---|---|---|---|---|
| No matches | No matches | No matches | No matches | No matches | No matches | No matches |

Total matches = 0

**Cistrome**

| Name | ID | Strand | Nb overlap columns | % aligned | Pearson correlation | Normalized cor |
|---|---|---|---|---|---|---|
| NAC_tnt.ANAC047_col_m1 | NAC_tnt.ANAC047_col_m1 | R | 14 | 0.8235 | 0.809 | 0.666 |
| NAC_tnt.NAM_colamp_a_m1 | NAC_tnt.NAM_colamp_a_m1 | D | 11 | 0.6875 | 0.878 | 0.604 |
| NAC_tnt.NAP_col_v3a_m1 | NAC_tnt.NAP_col_v3a_m1 | D | 11 | 0.6875 | 0.874 | 0.601 |

Total matches = 41 (38 more)

**Supplementary Figure S6.** *NAC61* regulative region analysis for *A. thaliana* ANAC047 and stress-related proteins *cis*-elements performed with the RSAT software. ANAC047, the NAC60 closest homolog, ORA47, RAP2.6 and RAP2.3, RRTF1, DEAR4, DREB2C, AP2EREBP, ABI3VP1 and G2like binding locations are reported.

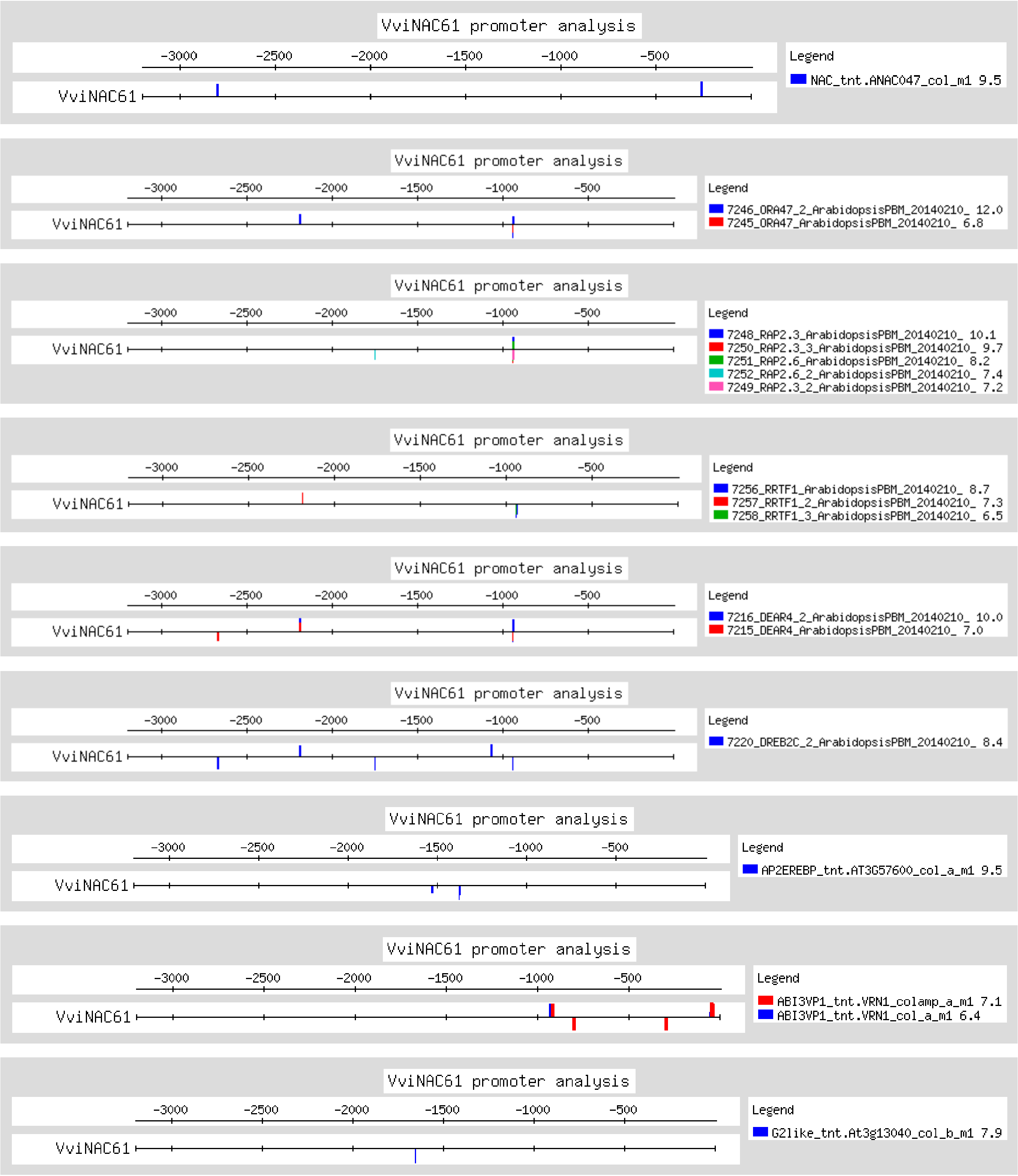

**Supplementary Figure S7.** Juice solute concentration in noble rot-induced and control grape berries of cv. 'Muller Thurgau'. (A) Sugar content (Brix°) and (B) Tritable Acidity trend throughout the post-harvest withering process; t1, t2 and t3 correspond to the three time points of grapes collection (Fig. 7B). Each value corresponds to the mean ± SD of three biological replicates.

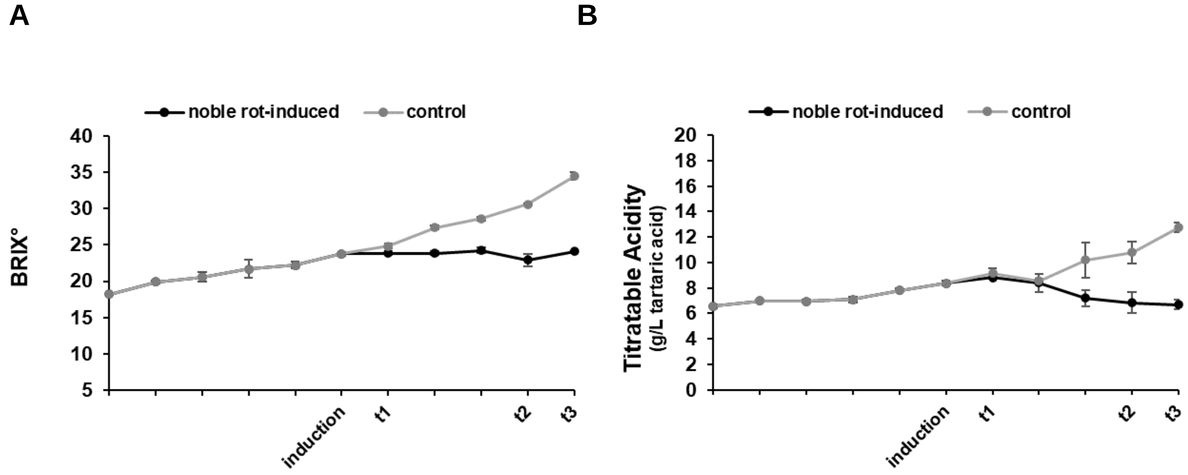

**Supplementary Table S1.** List of used primers.

| Gene ID | Functional annotation | Type of amplification | | Primers (5'-3') | Ta (°C) | Amplicon (bp) |
|---|---|---|---|---|---|---|
| VIT_08s0007g07640 | ***NAC61*** | CDS isolation | For | CACCATGGAAGAGGCTTCACTTG | 55 | 1212 |
| | | | Rev | TCAGTAGTCGAGTAAATAATCCA | | |
| | | Promoter isolation | For | CACCAATCTCTCTGAAAGCGGGGC | 53 | 1012 |
| | | | Rev | GGTTACACCACTACTGTATATCA | | |
| | | RT-qPCR | For | CGACCAGTACCGTAAGAGCC | 55 | 93 |
| | | | Rev | GCGAGAGGCTGACCATAGAC | | |
| VIT_07s0005g03340 | ***MYB14*** | Promoter isolation | For | CACCGGCTTCACCAATCATAGAGCTTA | 55 | 1569 |
| | | | Rev | TTTTTCTTTTCTACGTAAGGA | | |
| | | RT-qPCR | For | GATGTGGGAAATGGGAAATG | 54 | 150 |
| | | | Rev | CATTTGGCTGAGTCTGTTCG | | |
| VIT_04s0023g02480 | ***DHN1b*** | Promoter isolation | For | CACCCCCAACCACTCCACTACCAG | 54 | 814 |
| | | | Rev | TGTGTTGAAACGATCGATGAAATTT | | |
| VIT_17s0000g01280 | ***WRKY52*** | Promoter isolation | For | CACCTTGGTACACCACAAACGCAC | 54 | 938 |
| | | | Rev | TAGAGAGACTGAGAGAGATTGAGATTA | | |
| | | RT-qPCR | For | GAGTGGTGGACACCCATATCA | 56 | 102 |
| | | | Rev | AGTGATCATATCACAAGATCCTCCA | | |
| VIT_18s0001g01280 | ***LACCASE*** | RT-qPCR | For | TCACAGTGATTGGACCCGAA | 57 | 188 |
| | | | Rev | AATCAGAGGCATTGGGGTCA | | |
| VIT_08s0007g03030 | ***UBIQUITIN1*** | RT-qPCR | For | TCTGAGGCTTCGTGGTGGTA | 55 | 100 |
| | | | Rev | AGGCGTGCATAACATTTGCG | | |

**Supplementary Dataset S1.** *NAC61* co-expressed genes. The GCNs were obtained separately by looking at a berry-specific, a leaf-specific and a tissue-independent (TI) dataset. Key berry ripening regulators and DEGs during post-harvest dehydration are indicated.

**Supplementary Dataset S2.** Transcriptomic analysis of *NAC61*-overexpressing and control cv. 'Thompson Seedless' leaves. In sheet 2 and sheet 3 the upregulated and downregulated DEGs are reported, respectively. In sheet 2 berry post-harvest dehydration markers (Zenoni *et al*., 2016) and genes belonging to the STS GRN (Pilati *et al*., 2023) are reported.

**Supplementary Dataset S3.** Gene category MapMan distribution and enrichment analysis of DEGs.

**Supplementary Dataset S4.** NAC61 DAP-seq bound genes. In sheet 2 the promoter peaks selection (-3000≥bp ≥+100) is reported.

**Supplementary Dataset S5.** List of defined HCTs and detail of genes grouped in Fig. 5A. Genes belonging to NAC61 co-expressed genes (**Supplementary Dataset S1**), STS GRN (Pilati *et al.*, 2021), NAC60 VHCT (D'Incà *et al.*, 2023), and PHW markers (Zenoni *et al.*, 2016) are indicated. In sheets from 2 to 4 the DAP-seq, DEGs and GCNs exclusive genes are respectively reported; the common genes between 'GCNs and DAP-seq' and 'GCNs and DEGs' are reported in sheet 5 and 6, respectively; the DAP-seq, DEGs and GCNs common genes are listed in sheet 7.
